# LRCP: identifying functionally related SNPs and genes from phenome-wide genetic-effect correlations using GWAS summary statistics

**DOI:** 10.64898/2026.11.02.763742

**Authors:** Lane Guolan Chen, Thuan Quoc Thach, Pak C Sham

## Abstract

Genetic correlation describes which traits share genetic causes. The transposed question, which variants act alike across the phenome, has no comparable estimator, although it points directly to shared genes, pathways and regulators. Correlations of GWAS Z scores between variants across many traits are easy to compute, but they are confounded by linkage disequilibrium (LD) and, for any pair of variants, by sample overlap and genetic correlation among the GWAS. We show that in 452 Pan-UK Biobank GWAS the effective number of independent traits is only 58, and that naive tests reject 51-70% of null pairs of distal variants. We introduce LRCP (likelihood and regression on cross-products of Z scores), which estimates the phenome-wide genetic-effect correlation *ρ* between SNPs by regressing Z-score cross-products on LD cross components, using per-SNP heritability enrichment from the companion method LRCQ and LDSC-derived plug-ins. For variants in different LD blocks the estimator has a closed form, and its standard errors account exactly for the dependence among GWAS. In simulations LRCP is unbiased (∣bias∣ ≤ 0.03), its 95% intervals cover 93-98%, and its type-I error is 5.0-5.7%, whereas naive correlations are attenuated by 41-64% and reject up to 51% of null pairs. In Pan-UK Biobank, LRCP recovers the expected signs among *PCSK9*, *LDLR* and *APOB* variants and estimates *^ρ* = 0.90 between the *TP53* polyadenylation variant rs78378222 and *MDM2* rs3730556, whose Z scores correlate at *r* = 0.67 across traits (conditional bootstrap *P* = 0.015), although this correlation lies along a shared body-size and red-cell trait axis. Single variant pairs are near the resolution of current biobanks, with precision limited by the number of genetically distinct traits. Genome-wide, among 1,799 strongly enriched LD clumps (1.61 million distal pairs), a conditional test finds 437 pairs at FDR 5%, where naive correlation flags 1.1 million. Simulating the whole screen under the null shows that this test is conservative and that the 386 strongest links (*P* < 10^−5^) lie beyond every one of 322 million simulated null statistics. Linked genes share GO biological processes 1.9 times more often than random pairs of enriched loci. The links form 14 modules that match known biology, among them platelet and red-cell production, lipoprotein metabolism (*PCSK9*, *LDLR*, *APOB*, *CETP*, *LPL*), myeloid differentiation (*IRF8*, *ITGA4*), bone density (*WNT16*), pigmentation (*HERC2*, *SLC45A2*), alkaline phosphatase (*ALPL*, *GPLD1*) and IL-6 signalling (*IL6R*, *CRP*). Most modules act within one trait domain, where LRCP shows shared trait profiles rather than a tested shared mechanism. Once LD and the dependence among GWAS are modelled, phenome-wide summary statistics can identify functionally related variants and genes.

## Introduction

Most trait-associated variants are pleiotropic. Across the GWAS Atlas, a typical associated locus is linked to traits in several domains [Watanabe 2019], and pleiotropy pervades the human genome [Solovieff 2013; Jordan 2019]. Pleiotropy is usually summarised from the trait side. The genetic correlation *r_g_*between two traits is the correlation of their causal effects across SNPs [Bulik-Sullivan 2015b], and local extensions place that correlation in a genomic region [Shi 2017; Lu 2017; Zhang 2021; Werme 2022]. These quantities tell us which *traits* share genetic causes. They do not say which *variants* act alike. Two variants that affect the same set of traits in the same proportions and directions are functionally similar, whether or not they sit near each other. Plausible reasons include regulating the same gene, acting on two genes of one pathway, or converging on a shared trans-regulated core gene, as posited by the omnigenic model [Boyle 2017; Liu 2019; Võsa 2021]. Fig. 1a shows the simplest case. A cis-eQTL of a transcription factor and a cis-eQTL of the factor’s trans-regulated target both act on downstream traits through the target’s expression. Their effects are therefore proportional across traits, although the two variants are unlinked.

**Fig. 1.**
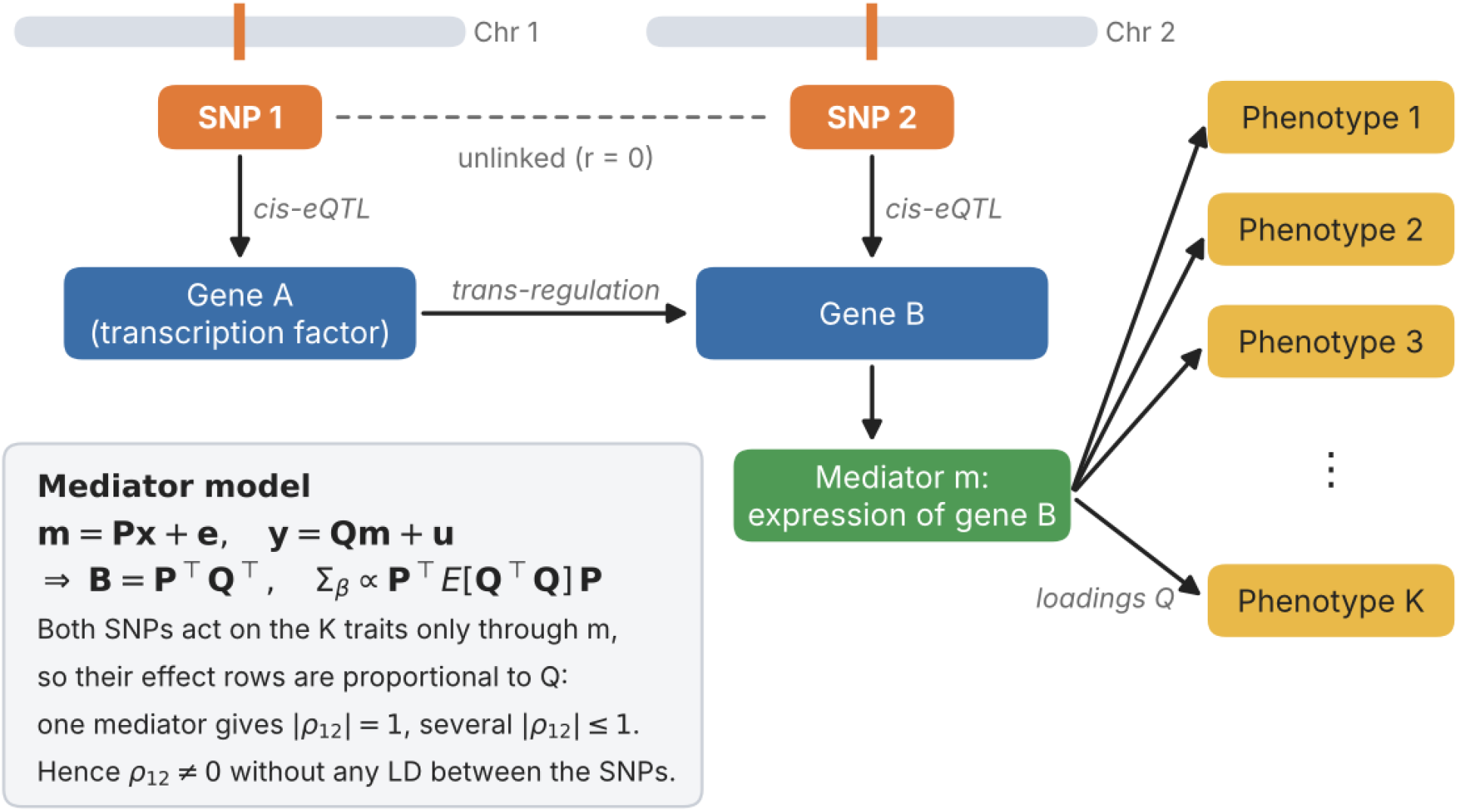
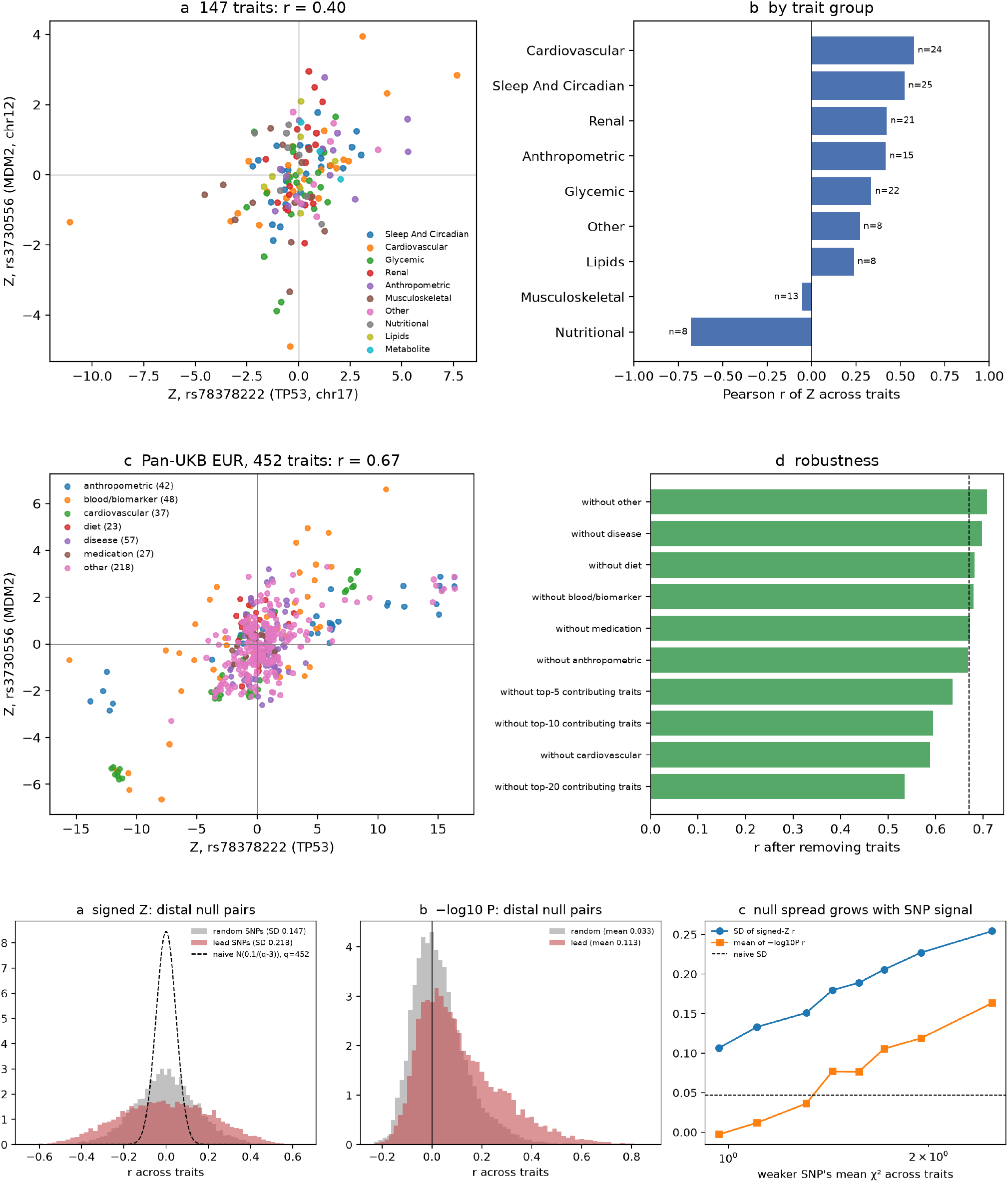
Phenome-wide genetic-effect correlation: mechanism, observation and null. (a) How two unlinked variants acquire a genetic-effect correlation: a cis-eQTL of a transcription factor (gene A) and a cis-eQTL of its trans-regulated target (gene B) act on *K* traits only through gene B’s expression, so under the mediator model **B** = **P**^⊤^**Q**^⊤^ their effect rows are proportional (Supplementary Note §S2.3). (b,c) TP53 rs78378222 × MDM2 rs3730556 in the CMDKP PheWAS, overall and by domain. (d,e) The same pair in 452 Pan-UKB traits, and after leaving out each trait domain or the most strongly associated traits. (f) Empirical null of 8,229 distal pairs vs naive and *q*_eff_ references.

We call this transposed quantity the **phenome-wide genetic-effect correlation** *ρ_ij_*. It is the correlation between the joint effects of SNPs *i* and *j* across a population of traits. *ρ_ij_* is to a pair of SNPs what *r_g_* is to a pair of traits. For a full effect matrix **B** (SNPs × traits), the correlation of its columns gives the genetic correlation matrix and the correlation of its rows gives the genetic-effect correlation matrix **R***_β_*. Large biobank GWAS collections with hundreds of heritable traits measured in one cohort [Bycroft 2018; Pan-UKB] now make the row-wise view estimable.

Hints of non-zero *ρ* are easy to find. Two variants on different chromosomes, rs78378222 in the polyadenylation signal of *TP53* [Stacey 2011] and rs3730556 in *MDM2*, the E3 ligase that degrades p53 in a negative feedback loop [Momand 1992; Bond 2004], have Z scores that correlate at *r* = 0.40 across 147 cardiometabolic, renal, sleep and anthropometric traits (Fig. 1b). The correlation is strongest among cardiovascular traits (*r* = 0.58) and survives removal of the most strongly associated traits (*r* = 0.37). It is seen again in a second resource, whose participants may overlap with those of the portal: across 452 Pan-UK Biobank traits the same two variants correlate at *r* = 0.67, with alleles that move pulse pressure, red-cell indices and lean mass in the same direction (Fig. 1d,e). Among gene-level association statistics across 4,756 GWAS ATLAS studies, the most correlated pairs between distal chromosomal segments include known functional partners: *PCSK9*–*APOB* in LDL metabolism, *SEC16B*–*TMEM18* in obesity, *IL23R*–*ATG16L1* in inflammatory bowel disease, and *TGFB2*–*LTBP1*–*EFEMP1* in TGF-β/extracellular-matrix biology (Supplementary Table S1).

Such observations cannot be taken at face value. GWAS summary statistics are *marginal* associations, so a SNP’s Z score carries the effects of every causal variant it tags through linkage disequilibrium (LD). Nearby SNPs therefore have correlated Z scores even when their causal effects are unrelated (“statistical link = functional link + LD”). Less appreciated is a second confounder that affects every pair, distal ones included: the GWAS are not independent. Biobank GWAS share participants, so their sampling errors correlate through phenotypic correlation [Bulik-Sullivan 2015b], and their true effects correlate through genetic correlation. We show that this dependence leaves the *mean* of a naive SNP-pair Z correlation at zero under the null but inflates its *variance*. Under realistic UK Biobank trait structure, a naive test of 300 traits has the precision of 16-120 independent ones and rejects 20-67% of truly null distal pairs. In the 452 Pan-UK Biobank traits the effective number of traits is only 58, and naive tests reject 51% of random and 70% of strongly pleiotropic null distal SNP pairs (Fig. 1f). Correlations of unsigned statistics such as − log_10_ *P* are also biased upwards (null mean *r* up to 0.26 with independent traits and 0.38 with genetically correlated traits) because traits differ in power. No current method separates genetic-effect correlation from these confounders. Factorisation and clustering of variant-by-trait association matrices [Tanigawa 2019; Udler 2018; Smith 2024; Ballard & O’Connor 2022] work on marginal statistics of lead variants and do not model LD or GWAS dependence. Colocalisation methods [Giambartolomei 2014; Foley 2021; Arvanitis 2022] ask whether one variant is causal for several traits, not whether two variants act alike. Gene-set methods correct gene-gene correlation for LD within one GWAS [de Leeuw 2015; Lamparter 2016] but not across GWAS.

Here we introduce **LRCP** (likelihood and regression on cross-products of Z scores). LRCP models phenome-wide GWAS Z scores under a random-effect model in which the SNP-side covariance of joint effects factorises into per-SNP enrichment *w* (estimated by our companion method LRCQ [Chen 2026]) and the genetic-effect correlation matrix **R***_β_*. We derive the expected cross-product of any two Z scores from any two GWAS as a sum of three components: shared sampling noise through LD, tagged effects, and genetic-effect correlation.

LRCP estimates **R***_β_* by regressing Z-score cross-products on LD cross components, with LDSC-derived intercepts and co-heritabilities as plug-ins. For SNPs in different LD blocks this reduces to a closed-form LD deconvolution of the trait-averaged cross-product matrix, restricted to candidate causal SNPs with non-zero enrichment. We derive standard errors that account for the full dependence among GWAS. In simulations, LRCP is approximately unbiased (∣bias∣ ≤ 0.03) and keeps type-I error at 5.0-5.7%, whereas naive correlations are attenuated by 41-64% through LD and reject up to 51% of null pairs when traits are correlated (Fig. 2). Applied to 444 Pan-UK Biobank traits with UK Biobank in-sample LD, LRCP recovers the expected signs among lipid-gene variants and estimates *^ρ* = 0.90 for TP53–MDM2, while showing that single SNP pairs are near the resolution limit of current biobanks (Fig. 3). Among 1,799 strongly enriched loci genome-wide, LRCP finds 437 distal pairs at FDR 5% (where naive correlation flags 1.1 million), grouped into modules of known biology such as platelet and red-cell production, lipoprotein metabolism, myeloid differentiation and IL-6 signalling. Linked genes share GO biological processes 1.9 times more often than random pairs of enriched loci (Fig. 4).

**Fig. 2.**
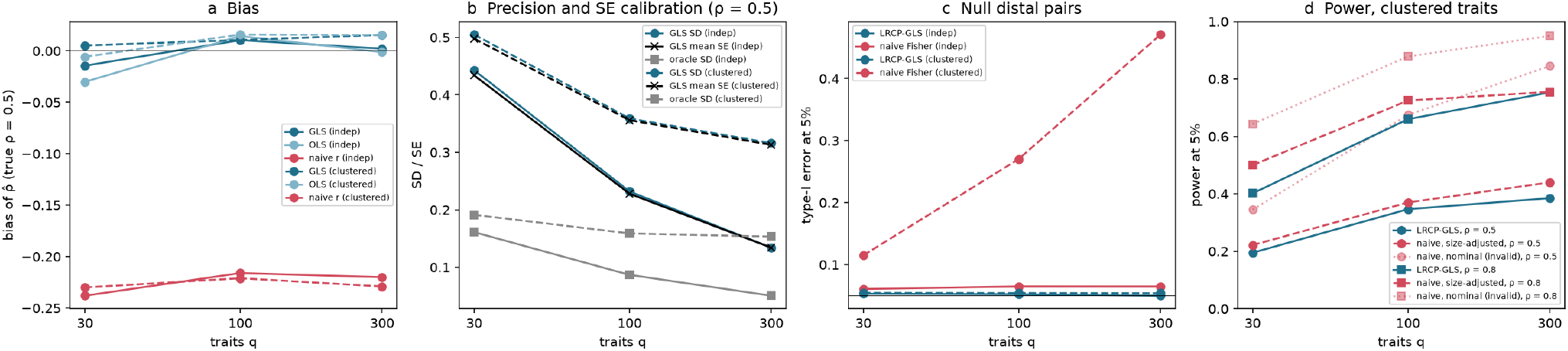
Simulations (S1). Two LD regions on different chromosomes, 200 replicates per cell, *q* ∈ {30, 100, 300} independent (solid) or clustered (dashed) traits, LD *r* = 0.6. (a) Bias of *^ρ* at *ρ* = 0.5 for LRCP-GLS, LRCP-OLS and the naive Z correlation. (b) Empirical SD of the GLS estimate, its mean analytic SE and the SD of the oracle estimator that uses the true effects. (c) Type-I error at the 5% level for null distal pairs, LRCP-GLS against the naive Fisher test. (d) Power at the 5% level for clustered traits: LRCP-GLS and the naive test after size adjustment (critical value from each test’s own null distribution), and the naive test at its nominal level, whose apparent power reflects its inflated type-I error.

**Fig. 3.**
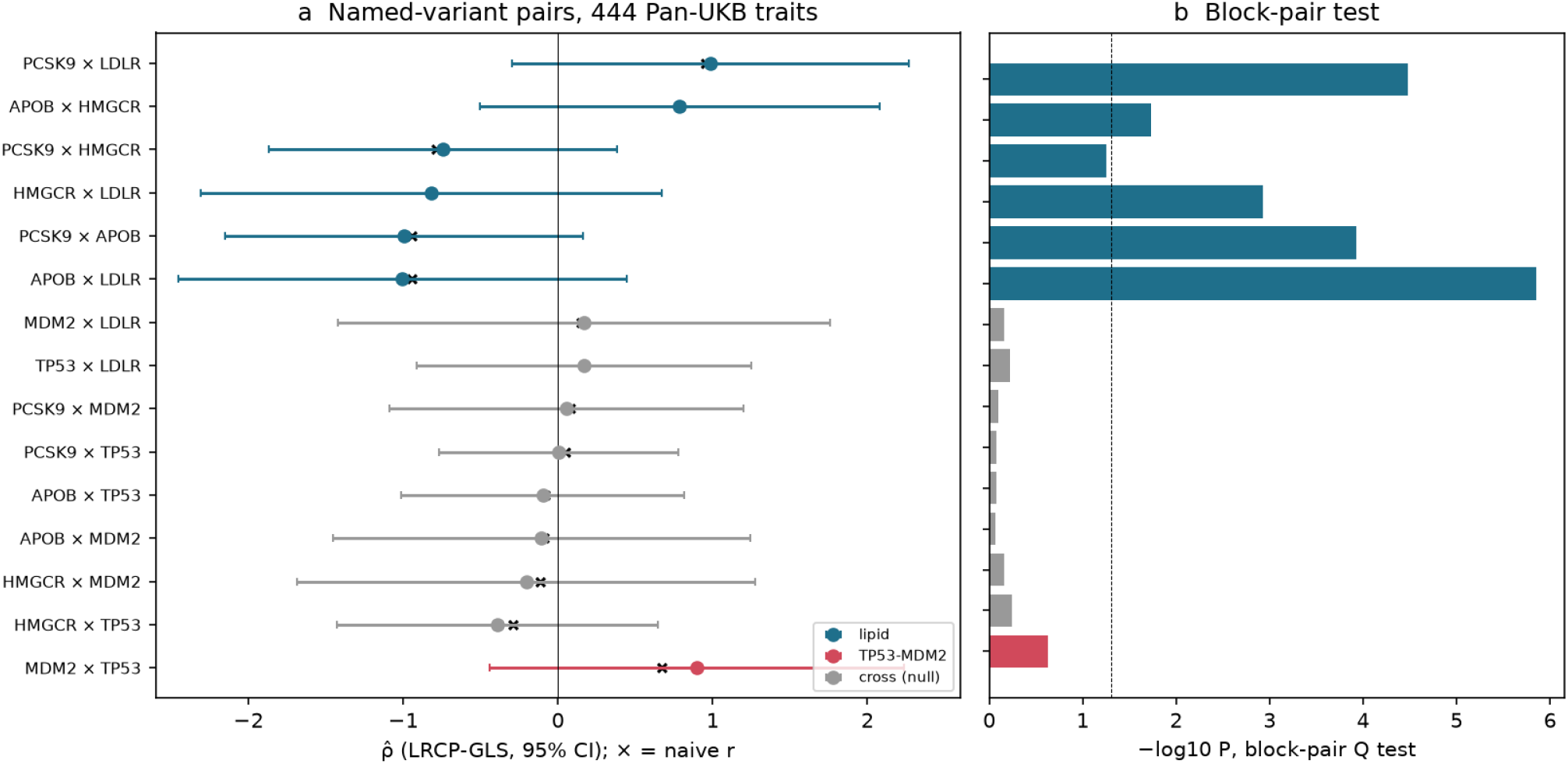
Targeted LRCP in Pan-UKB (444 traits, UK Biobank in-sample LD). (a) GLS *^ρ* with 95% CIs (plug-in SE) and naive *r* for the 15 named-variant pairs; (b) block-pair Q test.

**Fig. 4.**
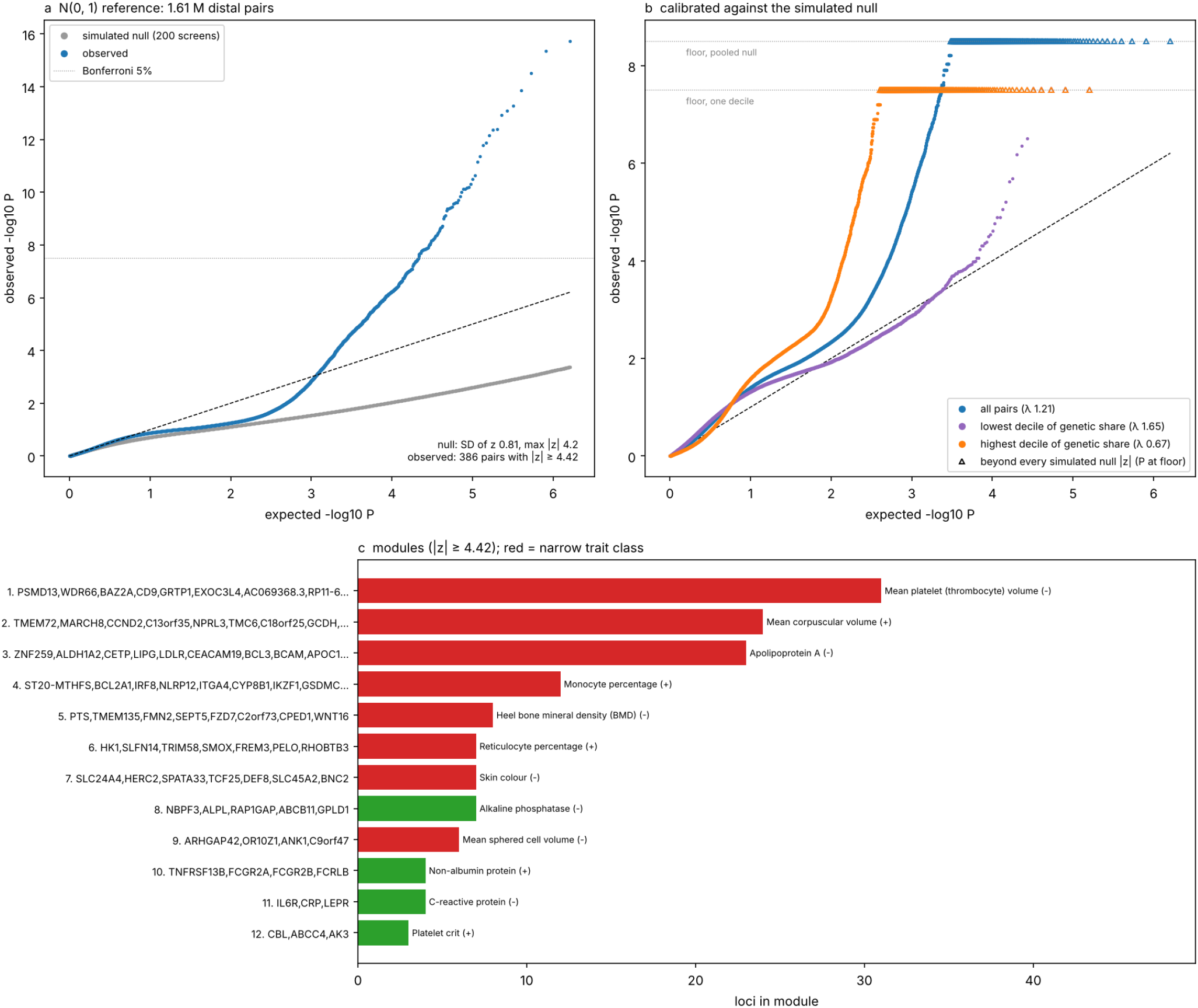
Genome-wide LRCP among 1,799 enriched tags (1.61 million distal pairs, 444 traits). (a) QQ plot of the conditional test against the *N* (0, 1) reference (blue), with the same statistic computed on 200 screens simulated under the separable null (grey; 322 million null pairs). The statistic is self-normalised, so its null is light-tailed (SD 0.81) and the normal reference is conservative; the observed bulk runs at or just below the diagonal, and the late rise is signal beyond the simulated null (386 observed pairs with ∣*z*∣ ≥ 4.42; the largest of 322 million null ∣*z*∣ is 4.22). Dotted line: Bonferroni 5% level. (b) QQ plot of P-values calibrated against the simulated null, for all pairs and for the deciles of pairs with the lowest and highest genetic share of their conditional variance (*λ*_cal_ 1.65 and 0.67; purple and orange). A calibrated P cannot be smaller than one over the number of simulated null statistics: pairs whose ∣*z*∣ exceeds every simulated null value (open triangles; 530 pairs overall, 400 in the highest decile) sit on that floor, 1/(*N* ^∗^ + 1) with *N* ^∗^ = 3.2 × 10^8^ for the pooled null and one tenth of that within a decile (dotted lines). (c) The 12 largest modules (connected components of pairs with *P* < 10^−5^), with each module’s top trait and its sign; red, at least one locus in a narrow trait class (descriptive, §S8.5); green, no locus in a narrow class.

## Results

### Phenome-wide Z-score correlations between distal SNPs: an observation and its null

#### Observation

rs78378222, in the polyadenylation signal of *TP53* on chromosome 17, and rs3730556 in *MDM2* on chromosome 12 are on different chromosomes and share no LD. Yet across 147 traits in the Common Metabolic Diseases Knowledge Portal (CMDKP) PheWAS, their meta-analysed Z scores correlate at *r* = 0.40 (Spearman 0.41) (Fig. 1b). The correlation is strongest for cardiovascular traits (*r* = 0.58, 24 traits), sleep and circadian traits (0.52) and renal and anthropometric traits (0.42). It remains 0.37 after removing the five traits most strongly associated with the *TP53* variant (Fig. 1c). The two variants are in linkage equilibrium in every LDpair reference panel (*r*^2^ < 0.005).

#### Replication in one cohort

In 452 Pan-UK Biobank European GWAS (phenotype QC pass, both sexes), the same two variants correlate at *r* = 0.67 (Fig. 1d). The rs78378222-G and rs3730556-G alleles both lower pulse pressure, mean corpuscular volume and haemoglobin, and both raise red-cell count and lean mass. The correlation is 0.95 among cardiovascular traits (37), 0.94 among anthropometric traits (42) and 0.72 among blood biomarkers (48). It is 0.59 without cardiovascular traits and 0.53 after dropping the 20 most contributing traits (Fig. 1e). In Pan-UKB’s maximal independent set of 150 traits, *r* = 0.41. The higher full-panel value does not conflict with the portal’s 0.40. Pan-UKB contains blocks of near-duplicate traits from one cohort (for example 37 cardiovascular and 42 anthropometric measures) on which the two variants align. These blocks raise *r* without adding independent evidence. Once the panel is pruned to independent traits, the two resources agree (0.41 vs 0.40). The null below accounts for exactly this redundancy. In FinnGen R12 [Kurki 2023], whose participants do not overlap with UK Biobank, the marginal Z scores of the same two G alleles correlate at *r* = 0.36 across 14 disease and medication endpoints, one per source-defined chapter chosen by case count alone (0.17-0.42 when any one endpoint is left out; Fig. S4). This is a descriptive comparison of marginal association profiles, with no LD correction and no null model; with 14 endpoints it is imprecise and is not an LRCP estimate.

#### How to judge such a correlation

Taken at face value with *q* − 3 degrees of freedom, *r* = 0.67 over 452 traits has *P* = 4 × 10^−66^. That reference is wrong. GWAS in one biobank share participants, and traits are genetically correlated, so the *q* Z scores of one SNP are far from independent. We show analytically (Supplementary Note §S9; simulations in Supplementary Table S9) that under the null of no genetic-effect correlation between two distal SNPs *k* and *l*, the naive correlation has mean zero but variance ≈ 1/*q*_eff_, with *q*_eff_ = tr **M***_k_* tr **M***_l_*/ tr(**M***_k_***M***_l_*) and **M***_k_* the trait-by-trait covariance of SNP *k*’s Z scores. *q*_eff_ shrinks with trait correlation, sample overlap and the SNP’s own pleiotropic signal.

In Pan-UKB the trait correlation of near-null SNPs gives *q*_eff_ = 58 for the 452 traits (70 for the independent set). We built an empirical null from 8,229 pairs of SNPs in 150 random 20-kb windows on different chromosomes or more than 5 Mb apart (Fig. 1f). The null SD of *r* is 0.147 for random SNPs and 0.218 for each window’s most pleiotropic SNP, against 0.047 implied by the naive test. **Naive Fisher tests reject 51% of random and 70% of lead-SNP null pairs at the 5% level** (16% and 22% in the independent set). Correlations of unsigned statistics such as − log_10_ *P* have positive null means (+0.03 for random and +0.11 for lead SNPs, +0.16 for the most pleiotropic pairs), because traits differ in power. This explains why gene-level − log_10_ *P* correlations across the GWAS Atlas look “ubiquitously” positive (Supplementary Table S1).

Against the matched empirical null, TP53–MDM2 remains unusual (*P*_emp_ ≈ 1 × 10^−4^, the resolution of thenull; *P* = 2 × 10^−9^ from *q*_eff_ ; independent set *P*_emp_ = 0.002). Lipid-gene controls (*LDLR* rs6511720, *PCSK9* rs11591147, *HMGCR* rs12916, *APOB* rs1367117) correlate at ∣*r*∣ = 0.79-0.95 with signs that follow their LDL-raising or LDL-lowering alleles. TP53 and MDM2 variants do not correlate with any lipid variant (∣*r*∣ ≤ 0.29, all *P*_emp_ > 0.25).

These calibrated marginal statistics still cannot separate genetic-effect correlation from the effects of other causal variants tagged through LD. That is what LRCP does.

### Overview of LRCP

LRCP models a SNP-by-trait matrix of joint effects as matrix normal, with SNP-side covariance proportional to **W**^1/2^**R***_β_***W**^1/2^ and trait-side covariance proportional to the genetic covariance matrix of the traits (Methods).

Here *w_i_* is the per-SNP enrichment of phenome-wide heritability estimated by the companion method LRCQ, and **R***_β_* holds the genetic-effect correlations *ρ_ij_*. The expected cross-product of the Z scores of SNPs *k* and *l* in GWAS *a* and *b* is the sum of three parts: shared sampling noise through LD (*c_ab_r_kl_*), tagged effects of the same causal SNPs (*t_ab_* ∑*_s_ w_s_r_ks_r_ls_*), and genetic-effect correlation between different causal SNPs (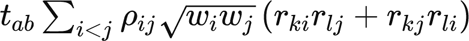). For SNPs in different LD blocks the first two parts vanish. LRCP then reduces to a closed-form deconvolution: average the cross-products over traits with weights from LDSC, and multiply on both sides by the inverse LD among the candidate causal SNPs (Supplementary Note §S4.6). Standard errors come from the exact Isserlis covariance of the cross-products under the full trait dependence. Candidates are prescreened on LRCQ enrichment, and correlations are reported at clump level (*r*^2^ < 0.5 tags) with variances re-estimated by the same LD projection.

### Simulations: LRCP is unbiased and calibrated where naive correlations fail

We simulated phenome-wide GWAS from the LRCP model with two LD regions on different chromosomes (2 × 2 AR(1) blocks of 100 SNPs; neighbour LD *r* ∈ {0.3, 0.6, 0.9}), 10% causal SNPs with log-normal enrichment, three cross-region causal pairs with *ρ* ∈ {0, 0.5, 0.8} (one negative), *q* ∈ {30, 100, 300} traits, and either independent traits or 10 trait clusters (within-cluster *r_g_* = 0.6, full sample overlap with phenotypic *r* = 0.3, as in UK Biobank). Each cell has 200 replicates and about 116,000 null candidate pairs (Methods, scenario S1).

#### Bias

LRCP-GLS is approximately unbiased in every cell (∣bias∣ ≤ 0.03 across 54 cells; OLS ≤ 0.05) (Fig. 2a). The naive Z correlation between the two causal SNPs is biased towards zero by 41-64% (about 45% at moderate LD: −0.22 at *ρ* = 0.5 and −0.37 at *ρ* = 0.8; about 60% at LD *r* = 0.9), because each causal SNP’s Z score also carries the effects of the other causal SNPs it tags.

#### Precision and SEs

With independent traits the SD of *^ρ* falls from 0.44 (*q* = 30) to 0.13 (*q* = 300) at *ρ* = 0.5. With clustered, overlapping traits it falls only from 0.51 to 0.32, because the effective number of traits is much smaller than *q* (Fig. 2b). The analytic SE matches the empirical SD within 10% except at LD *r* = 0.9 with *q* = 30, where it is up to 23% small. The null SE is exact at *ρ* = 0 and slightly small at large *ρ* (0.29 vs 0.36 at *ρ* = 0.8, *q* = 300, clustered). The plug-in SE that adds the second Isserlis term at *^ρ* removes this gap (0.34), and 95% intervals cover 93-98%.

#### Type-I error

Over about 2.1 million null pairs, LRCP-GLS rejects 5.0-5.7% at the 5% level in every cell (Fig. 2c). The naive Fisher test rejects 6-8% with independent traits and 11-13%, 26-30% and 47-51% at *q* = 30, 100 and 300 with clustered traits. Its error grows with *q* because correlated traits add precision to the naive SE but not to the estimate. In the bulk the GLS test is slightly conservative (genomic-control *λ* 0.84-0.98), while the naive test has *λ* = 1.5, 3.2 and 7.4 at *q* = 30, 100 and 300 with clustered traits (Supplementary Fig. S1).

In the far tail LRCP is inflated: at *P* < 10^−3^ it rejects 0.0012-0.0032 with independent traits and 0.0032-0.0040 with clustered traits, and at *P* < 10^−4^ up to 1.1 × 10^−3^. The cross-product statistic is a sum of products of Gaussians, so its tails are heavier than normal when the effective number of traits is small. The conditional test of Methods, which is exactly Gaussian given one window’s Z scores, removes this tail inflation. On the same 150 replicates (*q* = 100 clustered traits, 58,558 null pairs), the GLS test with model SEs rejects 5.2% at 5%, 0.31% at 10^−3^ (184 pairs against 59 expected) and 0.074% at 10^−4^ (44 against 6). The conditional test rejects 3.9%, 0 and 0. The conditional test is therefore the primary test for LRCP, and the GLS form is used for estimation.

#### Power

For a fair comparison we size-adjusted both tests, taking each test’s critical value from the 95th percentile of its own null ∣*z*∣ in the same cell (Fig. 2d). With independent traits and LD *r* = 0.6, the size-adjusted naive test is somewhat more powerful at small *q* (0.32 against 0.23 at *q* = 30 and 0.75 against 0.67 at *q* = 100, *ρ* = 0.5), and the two are equal at *q* = 300 (0.96 and 0.95; Table 2). With clustered traits, power is limited by the small effective number of traits for both: at *q* = 300 and LD *r* = 0.6, LRCP has power 0.75 at *ρ* = 0.8 and 0.38 at *ρ* = 0.5, against 0.76 and 0.44 for the size-adjusted naive test. LRCP gains in tight LD (LD *r* = 0.9: 0.60 against 0.48 at *ρ* = 0.8), where the naive correlation is diluted by the effects of tagged neighbours. The naive test’s apparently higher power at its nominal level (0.95 at *ρ* = 0.8) comes entirely from its inflated type-I error, and size adjustment needs the true null distribution, which is not available in real data.

**Table 1.** Methods that relate variants or traits through multi-trait GWAS summary statistics.

| Method | Estimand | Accounts for LD | Accounts for dependence among GWAS | Output and test |
|---|---|---|---|---|
| Naive Z-score correlation across traits (Fisher test) | Marginal association similarity of two SNPs | No | No (assumes independent traits) | *r*; invalid test (Fig. 1f) |
| Correlation of $-\log_{10} P$ or $\chi^2$ across traits | Similarity of association strength | No | No | Biased upwards under the null |
| LDSC, cross-trait LDSC, HDL [Bulik-Sullivan 2015a,b; Ning 2020] | *h*<sup>2</sup>, *r*<sub>*g*</sub> between two traits across SNPs | Yes (LD scores, full LD) | Yes (intercepts) | Genome-wide trait-pair *r*<sub>*g*</sub> |
| ρ-HESS, GNOVA/SUPERGNOVA, LAVA [Shi 2017; Lu 2017; Zhang 2021; Werme 2022] | Local *r*<sub>*g*</sub> between two traits in a region | Yes | Yes (sample overlap) | Trait-pair correlation per region |
| DeGAs, bNMF clustering, PDR [Tanigawa 2019; Udler 2018; Smith 2024; Ballard & O’Connor 2022] | Components or clusters of variant-by-trait association profiles | No (lead or pruned variants) | Partly or heuristically | Variant groupings; no pair-level test |
| coloc, HyPrColoc, CAFEH [Giambartolomei 2014; Foley 2021; Arvanitis 2022] | Whether one variant is causal for several traits | Within a locus | Partly | Posterior probability of a shared causal variant |
| MAGMA, Pascal [de Leeuw 2015; Lamparter 2016] | Gene and gene-set association in one GWAS | Yes, within one GWAS | Not across GWAS | Gene and pathway P-values |
| **LRCP (this work)** | **Genetic-effect correlation *ρ*<sub>*ij*</sub> of two SNPs or genes across a population of traits** | **Yes (deconvolution of the candidates’ LD)** | **Yes (intercepts and genetic covariances of all trait pairs)** | **$\hat{\rho}$ with SEs; conditional test exact given one window; gene, module and genome-wide tests** |

**Table 2.** Simulation S1 at LD. *r* = 0.6 **and** *ρ* = 0.5. Bias, empirical SD, mean plug-in SE and 95% interval coverage of LRCP-GLS; bias of the naive Z correlation; type-I error at the 5% level for null distal pairs; size-adjusted power at the 5% level. 200 replicates per cell; all cells are in Supplementary Table S3.

| **Traits** | ***q*** | **GLS bias** | **GLS SD** | **GLS mean SE** | **95% CI coverage** | **Naive *r* bias** | **Type-I, GLS** | **Type-I, naive** | **Power, GLS** | **Power, naive (size-adjusted)** |
|---|---|---|---|---|---|---|---|---|---|---|
| independent | 30 | -0.014 | 0.44 | 0.43 | 0.97 | -0.24 | 0.053 | 0.061 | 0.23 | 0.32 |
| independent | 100 | +0.010 | 0.23 | 0.23 | 0.96 | -0.22 | 0.052 | 0.065 | 0.67 | 0.75 |
| independent | 300 | +0.002 | 0.13 | 0.13 | 0.96 | -0.22 | 0.050 | 0.065 | 0.96 | 0.95 |
| clustered | 30 | +0.005 | 0.51 | 0.50 | 0.97 | -0.23 | 0.054 | 0.115 | 0.20 | 0.22 |
| clustered | 100 | +0.011 | 0.36 | 0.36 | 0.97 | -0.22 | 0.054 | 0.270 | 0.35 | 0.37 |
| clustered | 300 | +0.016 | 0.32 | 0.31 | 0.96 | -0.23 | 0.054 | 0.471 | 0.39 | 0.44 |

The advantage of LRCP is therefore validity and unbiased estimation rather than raw power; power for single SNP pairs is set by the number of genetically distinct traits (see *Interpretation* in the targeted analysis and §S9).

#### Prescreening on estimated enrichment (S4)

Replacing the true *w* by LRCQ estimates (*q* = 300 clustered traits, *ρ* = 0.5, 300 replicates) keeps LRCP unbiased for the true pairs that pass the screen (mean *^ρ* = 0.497-0.499) and conservative for null pairs (type-I error 3.3-3.8%). At thresholds *t* = 0.5-2 the screen keeps 18 candidates per region and 2.0 of the 3 true pairs. A stricter *t* = 4 keeps 13-14 candidates and 1.6 true pairs, and it raises BH discoveries from 0.03 to 0.05 per replicate (false discovery proportion 0.11 and 0.08). Discovery of single pairs at FDR 5% is rare at this signal level, which again points to aggregation as the route to discovery.

#### Non-separable effects (S7a)

We added to the null S1 background one SNP per region whose effects fall only on the traits of a single cluster (within-cluster effect correlation 0.8), drawn independently for the two regions, so the population *ρ* is 0. When the two loci act on different clusters, both the GLS and the conditional test stay calibrated (type-I error 2.0-4.0%). When they act on the same narrow cluster, both reject far too often: 9-11% for moderate locus effects (per-trait *Z* ≈ 3) and 28-31% for strong ones (*Z* ≈ 6), with null *z* SDs of 1.2-1.8. Two loci confined to one trait domain have only a few effective traits between them, so their realised correlation over the observed traits is large by chance, and a test against the superpopulation *ρ* = 0 that assumes the genome-wide trait covariance is anti-conservative. The two tests fail equally, so non-separability alone does not explain their disagreement for the lipid loci, but it does mean that correlations between loci acting on one narrow domain (such as the lipid genes) should be read as fixed-trait, within-domain similarity. The class null used in the real-data analyses below (Supplementary Note §S8.5) is our remedy: it takes the trait covariance of the test from other loci of the same trait-profile class.

#### Real LD and LD-reference mismatch (S5)

With UK Biobank in-sample LD for the *TP53* and *MDM2* blocks (464 and 962 HapMap3 SNPs), causal SNPs drawn from *r*^2^ < 0.5 tags (about 15 and 31 per block), and *q* = 100 or 300 clustered traits (100 replicates per cell), LRCP-GLS is unbiased (*^ρ* = 0.517 at *ρ* = 0.5) and the GLS null test rejects 5.2-5.8% of null pairs. The conditional test rejects 3.6-4.1% and is more powerful for the true pairs (0.55-0.66 vs 0.43-0.56). When the same data are analysed with 1000 Genomes EUR LD (503 samples; mean ∣Δ*r*∣ = 0.04 against UK Biobank LD), the estimate is nearly unbiased (0.53-0.55) but 20-25% more variable, and the GLS null test’s type-I error doubles to 10.7-11.3%. The conditional test stays calibrated (3.8-4.0%) with similar power (0.44-0.53). The conditional test is therefore also the robust choice when only an out-of-sample LD reference is available.

#### Plug-in errors (S6)

Errors in stage 1-2 inputs matter little except for genetic correlation. With 20% log-normal error in each trait’s *h*^2^, *^ρ* is biased by about −6% (0.48 vs 0.51 at *ρ* = 0.5) and both tests stay calibrated (4.0-4.5%). Errors of ±0.05 in the off-diagonal intercepts change nothing visible. Ignoring genetic correlation between traits (diagonal genetic covariance) leaves *^ρ* unbiased but inflates type-I error to 17% and 32% (GLS) and 13% and 23% (conditional) at *q* = 100 and 300 (150 replicates per cell). Genome-wide cross-trait LDSC *r_g_* is therefore a required input.

#### Gene-level aggregation (S9)

We placed two “genes” of three causal SNPs each in the two regions, with within-gene effect correlation 0.6 and a common cross-gene correlation *ρ* for all nine SNP pairs (*q* = 100 or 300 clustered traits, 150 replicates). The gene burden test is far more powerful than the best single SNP pair with Bonferroni correction over the nine pairs: 0.79-0.89 vs 0.45-0.66 at *ρ* = 0.5, and 0.35-0.37 vs 0.07-0.10 at *ρ* = 0.3. It is calibrated (6.0%) only when its variance includes the within-gene genetic covariance. With the default variance, which assumes no local correlation between a gene’s causal SNPs, it rejects 16-17% of null gene pairs, because correlated effects within a gene make the burden more variable. Gene-level LRCP therefore needs an estimate of the local genetic covariance. The local moment 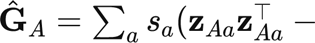 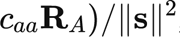, with P-values from a conditional parametric bootstrap that redraws **Z***_A_* with **Z***_B_* fixed, is the package default (§S8.2a). Repeating S9 with it (100 replicates per cell) gives type-I error 4-6% and power 0.90-0.92 at *ρ* = 0.5, 0.43-0.48 at *ρ* = 0.3 and 0.13-0.25 at *ρ* = 0.2. Over the nine SNP pairs with Bonferroni correction, the best single pair has power 0.66-0.79, 0.24-0.28 and 0.03-0.19. The default test even exceeds the oracle-covariance version (0.75-0.86 at *ρ* = 0.5), because its variance is estimated from the realised local signal.

#### Local pairs (S2)

Inside one LD block (AR(1), neighbour *r* = 0.9; *q* = 300 clustered traits; 200 replicates), the naive Z correlation of two causal SNPs mostly reflects their LD. With *ρ* = 0 its median is 0.43, 0.75 and 0.96 for LD *r* = 0.21, 0.48 and 0.81. LRCP-local removes the LD and tagging components and is close to unbiased: the median *^ρ* is −0.07 to 0.04 at *ρ* = 0, and 0.38-0.47 at *ρ* = 0.5, with mild attenuation at *r* = 0.81. It is imprecise, though (MAD 0.35-0.50, against SDs of 0.13-0.19 for comparable distal pairs at *q* = 300), because the correlation component must be separated from much larger LD terms. We therefore concentrate on distal pairs.

#### A small design and the pair-counting convention (S8)

In a small design (100 SNPs in two blocks of 50, 200 GWAS, one causal pair), a regression of Z cross-products on LD cross components that counts each SNP pair twice is attenuated and unstable: its slope is about *ρ*/2 (pooled slope 0.53 at *ρ* = 0.9). A Gaussian likelihood is almost as accurate as using the true effects (RMSE/∣*ρ*∣ 2.5%, 11% and 36% at ∣*ρ*∣ = 0.9, 0.5 and 0.2, against 69-440% for OLS/WLS and 85-94% for naive Z correlation). Counting each unordered pair once (Supplementary Note §S1.5) removes the halving: the regression slopes are 0.817, 0.485, 0.205 and −0.518 for true *ρ* = 0.8, 0.5, 0.2 and −0.5 (Supplementary Table S6).

#### Other misspecified effects (S7b-c)

We added one locus per region to the S1 background, with effects that violate the matrix-normal model in two ways (200 replicates per cell).

- *Sparse effects (S7b).* Each locus acts on its own random 10% of traits, independently of the other (true *ρ* = 0). Both tests stay conservative (type-I error 0-2.5%).
- *Shared mediator, same trait (S7c).* Both loci act through the same mediator trait, so their effects follow that trait’s genetic-correlation profile (∣*ρ*∣ = 1). LRCP recovers the correct sign with *^ρ* = 0.76-0.92 (shrunk by background polygenic signal) and rejects in 94-100% of replicates.
- *Shared mediator, different traits (S7c).* Each locus has its own mediator (mean ∣*ρ*∣ 0.16-0.23). *^ρ* is attenuated (0.04-0.11) and the tests reject 6.5-15% of the time.

Mediated pleiotropy through a shared trait is therefore detected as a strong genetic-effect correlation, which is the pattern seen for the lipid and blood-cell modules.

#### Likelihood versus GLS (S1-MLE)

In the small design a Gaussian likelihood was far more accurate than regression (S8 above). We compared lrcp_mle with the GLS estimator in the S1 design at *q* = 30 clustered traits and LD *r* = 0.6: 10 replicates of three causal pairs each at *ρ* = 0 and at ∣*ρ*∣ = 0.5. The MLE was fitted on a screened set of the three causal SNPs plus two other candidates per region, and GLS on all candidates. The MLE was markedly more precise, with RMSE 0.30 against 0.50 for GLS at *ρ* = 0 and 0.32 against 0.63 at ∣*ρ*∣ = 0.5. Both had small bias (MLE −0.01 and −0.07, GLS −0.07 and +0.05). MLE estimates stayed within [−1, 1], whereas 3-17% of GLS estimates fell outside. This agrees with the small design and with the numerical check in Supplementary Note §S4.5, in which the exact bounded likelihood also handled SNPs with zero enrichment (*R*^2^ 0.905 against 0.854 for OLS). Two findings keep GLS as the default. First, the MLE’s Hessian standard errors are too small: its 95% intervals covered the truth in 63-70% of cases and its Wald test rejected 30% of null pairs, whereas the GLS plug-in intervals covered 97%. Second, the MLE took several minutes per replicate for five SNPs per region, against seconds for GLS on all candidates, and part of its advantage here comes from the small candidate set that contains the true pairs. We therefore use the MLE to refine point estimates within small screened sets and base all tests on GLS and the conditional test.

### LRCP in Pan-UK Biobank: targeted analysis of TP53–MDM2 and lipid genes

We applied LRCP to the six LD blocks (Berisa-Pickrell EUR) containing the TP53, MDM2, PCSK9, LDLR, HMGCR and APOB variants: 4,788 HapMap3 SNPs plus the two named non-HapMap3 variants, 444 traits, and UK Biobank in-sample LD (Weissbrod et al. 2020). Stage 1-2 plug-ins are the genome-wide cross-trait LDSC matrices of the 444 traits used by LRCQ, with *n* the total sample size of each GWAS (Methods). Sensitivity analyses with approximate plug-ins (Cheverud *r_g_* ≈ *r_p_*) or with the effective sample size of binary traits gave nearly identical *^ρ*.

#### Enrichment

Clump-level LRCQ enrichment is highest near the lipid genes (for example *LDLR* tag 19:11210912 *^w* = 1,214, *z* = 4.1; *PCSK9* rs11591147 *^w* = 946, *z* = 2.8; *APOB* region tags *^w* = 128-301, *z* = 8-20) and in the *TP53* block (several tags with *^w* = 82-98, *z* = 6.5-10). The *TP53* polyadenylation variant itself is rare (allele frequency 1.3%) and its enrichment is imprecise (*^w* = 321, SE 744).

#### Stage 4

With candidates defined as tags with enrichment *z* ≥ 3 plus the named variant in each block (2-16 per block, 977 cross-block pairs), the GLS estimate is *^ρ* = 0.90 (SE 0.59, *P* = 0.13) for TP53 rs78378222 × MDM2 rs3730556. The lipid controls have the signs expected from their LDL effects. PCSK9 rs11591147 × APOB rs1367117 gives *^ρ* = −0.99 (SE 0.46, *P* = 0.03), PCSK9 × LDLR rs6511720 0.99 (SE 0.54, *P* = 0.07), APOB × LDLR −1.01 and APOB × HMGCR 0.79. TP53 and MDM2 variants against lipid variants give ∣*^ρ*∣ ≤ 0.39 with *P* ≥ 0.44. No single SNP pair passes BH correction across the 977 pairs.

#### Block-pair tests

Under the separable-model conditional null (Methods), the orientation-free statistic *Q* aggregated over all candidate pairs of a block pair separates the lipid blocks: APOB-LDLR *P* = 1 × 10^−6^, PCSK9-LDLR 3 × 10^−5^, PCSK9-APOB 1 × 10^−4^ and HMGCR-LDLR 0.001. The bootstrap max-T over tag pairs gives *P* ≤ 0.001 for the first three and *P* = 0.05 for HMGCR-LDLR. Every block pair involving TP53 or MDM2 has *P* ≥ 0.23 (TP53-MDM2 *P* = 0.23). Every lipid-variant estimate has the sign predicted by the variants’ effects on LDL cholesterol (PCSK9 rs11591147-T and LDLR rs6511720-T lower LDL; APOB rs1367117-A and HMGCR rs12916-C raise it).

#### Inference for loci confined to a narrow trait domain

For the lipid block pairs the conditional test gives *z* about 2.8 times larger than the GLS test of the same block-level statistic (PCSK9 × APOB −7.3 vs −2.6; PCSK9 × LDLR 6.4 vs 2.6; variance ratio *R_AB_*0.12-0.16). The two are equal in expectation under the model and are calibrated and equally powerful in simulations under it, so the gap signals a departure from the model. Two departures matter (Supplementary Note §S8.5). First, the GLS denominator uses the side-B enrichment *^w_B_*, which is noisy and clipped (rs78378222 *^w* = 321 ± 744), while the conditional test uses the realised Z profile of side B. Second, and more important, loci of one functional class share a narrow trait profile: the signal of each target block spans only 2-5 effective trait dimensions (tr(**S**)^2^/ tr(**S**^2^) for the trait covariance **S** of the block’s enriched tags after subtracting the intercepts), against *q_G_* ≈ 11 for the genome-wide genetic covariance that the plug-in variance uses. Independent loci of such a class have nearly collinear trait profiles, so their realised correlation over the observed traits is large even when the population *ρ* is 0 (simulation S7a). The remedy is a class null, in which the trait covariance of the test variance comes from other loci of the same class, held out from the pair being tested (§S8.5). When the class has fewer than about five effective traits, as for the lipid loci, no test of alignment against that null has power. We therefore report the lipid *^ρ* ≈ ±1 descriptively, as the signature of a shared LDL-cholesterol axis with biologically correct signs, not as significant correlations.

#### Conditional test for TP53 × MDM2

The conditional test with the local-moment variance and bootstrap P-values (larger P of the two orientations) gives *P* ≤ 0.001 for all six lipid pairs, *P* ≥ 0.48 for every TP53/MDM2 × lipid pair, and *P* = 0.015 for TP53 × MDM2 (GLS *P* = 0.13). Conditioning on MDM2 gives *z* = 2.59, and conditioning on TP53 gives *z* = 1.67 (normal *P* = 0.010 and 0.095; bootstrap *P* = 0.011 and 0.015). The statistic is self-normalised: the trait covariance of the conditioned locus is re-estimated from its own Z profile. Its null distribution is therefore light-tailed, and the normal reference is conservative. In simulations at this configuration the bootstrap is calibrated (3.3-4.5% at 5%, 0.7% at 1%), while the normal reference rejected nothing. We report the bootstrap P from the less significant orientation, *P* = 0.015. This assumes the genome-wide trait-covariance model. The shared signal lies in red-cell indices and body size, so a shared-mediator reading (§S8.5) cannot be excluded. LRCQ’s TP53 gene-level enrichment is driven mostly by the nearby *SHBG* cis-signal, but SHBG contributes only 2.6% of rs78378222’s phenome-wide *χ*^2^. Dropping it leaves the test unchanged (*z* 2.61 and 1.67, bootstrap *P* ≤ 0.012). Under the class null (§S8.5), each variant’s class consists of the genome-wide candidate tags whose deconvolved trait profiles correlate with it at ∣*r*∣ ≥ 0.5 . Both classes span fewer than five effective traits (*q*_eff_ 2.5 for MDM2 and 2.3 for TP53), and the class-null test is not significant (*z* = 1.35, *P* = 0.18; at ∣*r*∣ ≥ 0.3, *q*_eff_ 4.2 and 3.7, *P* = 0.26). The TP53–MDM2 correlation therefore lies along a shared body-size and red-cell trait axis, and current data cannot separate it from shared membership of that axis.

#### Interpretation

The point estimates agree in sign and size with the calibrated marginal analysis, but single SNP pairs are near the resolution limit of 444 UK Biobank traits. Strong enrichment does not rescue them. For large *w* the variance of *^Cij* is dominated by its genetic term *s*^2^*w_i_w_j_*/*q_G_* (Supplementary Note eq. 9.1), so SE(*^ρ_ij_*) approaches a floor of about 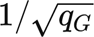 set by the number of genetically distinct traits, whatever the SNPs’ enrichment. The genome-wide genetic trait covariance of the 444 traits has *q_G_* ≈ 11 effective traits 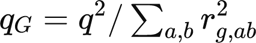 over the genome-wide LDSC genetic correlations, Methods; the sampling-noise correlation of near-null SNPs gives a much larger 58), so the floor is about 0.30. Deconvolving 2-16 candidates in partial LD (*r*^2^ < 0.5) inflates it to the observed 0.40-0.90. Aggregation across the candidate SNPs of a gene or block pair (gene-level test, Methods) is therefore the natural unit for discovery, and adding genetically distinct trait domains, not more GWAS of correlated traits, is what raises power.

### Genome-wide LRCP among enriched SNPs

We took the 1,799 LD-clump tags that LRCQ finds enriched for phenome-wide heritability (score-type test, *q* < 0.05 for *W_t_*> 0; in 868 of the 1,383 3-Mb windows; 444 traits; companion paper) and tested every distal pair (different chromosomes or ≥ 5 Mb apart; 1.61 million pairs) with the conditional test, conditioning on the other enriched tags of each window (LD among them from 1000 Genomes EUR, max *r*^2^ 0.49; plug-ins from genome-wide cross-trait LDSC). Naive Z-score correlation on the same pairs rejects 73% of pairs at *P* < 0.05 and 651,470 at Bonferroni level, as expected from the shared samples and genetic correlations among the traits.

#### Null calibration of the screen

Against the *N* (0, 1) reference the QQ plot of the 1.61 million conditional *z* runs at or just below the diagonal in the bulk and then rises steeply across it (Fig. 4a): 0.82% of pairs have *P* < 0.05, yet 386 pairs have *P* < 10^−5^ (∣*z*∣ ≥ 4.42). The bulk lies low because of how the statistic is built, not because of the data. The test variance uses the local-moment trait covariance *^GA*, estimated from the same Z profile as the numerator, and the larger of the two orientation variances, so the statistic is self-normalised and light-tailed under the null (as for TP53 × MDM2 above). To show this directly we simulated the whole screen under the fitted separable null: every window was redrawn as **Z**^∗^ ∼ *N* (0, Γ ⊗ *R_W_* + *T* ⊗ *^GW*), with *^_GW_* the PSD-projected local moment, and the screen, including *^G^∗^*, was recomputed (200 replicates, 322 million null pairs; Methods). The simulated null is lighter-tailed still: its *z* has SD 0.81 (observed 0.93), 0.36% of null pairs have *P* < 0.05, and the largest ∣*z*∣ among 322 million null pairs is 4.22 (Fig. 4a, grey). Beyond ∣*z*∣ ≈ 3 the observed and null distributions separate, with 46 times as many observed pairs as null pairs at ∣*z*∣ 3-3.5, 429 times as many at 3.5-4 and about 12,000 times as many at 4-5; 158 observed pairs exceed ∣*z*∣ = 5. No null pair in any of the 200 simulated screens reached ∣*z*∣ = 4.42 (calibrated *P* ≈ 3.1 × 10^−9^ at that threshold, the resolution of the pooled null). If the screens are independent, the probability that a null screen has at least one pair beyond this threshold is below 1.5% (one-sided 95% bound), against 386 observed pairs. The normal reference is therefore conservative, and the upper tail is real signal relative to the separable null.

Calibrated against the simulated null (Fig. 4b), the genomic-control factor of the calibrated P-values is *λ*_cal_ = 1.21 (median of the corresponding *χ*^2^ statistics over its null value). How the bulk departs from the null depends on how much of each pair’s projected variance is genetic: *λ*_cal_ falls from 1.65 in the decile of pairs with the lowest genetic share to 0.67 in the decile with the highest (Table S7). The highest decile has a deflated bulk, yet it holds most of the extreme tail: of the 530 pairs whose ∣*z*∣ exceeds every simulated null value, 400 are in that decile and 5 in the lowest. This is the pattern expected when enriched loci concentrate on a few shared trait axes (blood-cell indices, lipids, bone density, pigmentation) instead of spreading their effects over the genome-wide genetic trait covariance *T* as the separable null assumes. Loci on different axes then have nearly orthogonal profiles, with smaller cross-products than the null, and loci on the same axis have nearly collinear profiles even without a specific shared mechanism (simulation S7a, §S8.5). With the calibrated reference 3,154 pairs pass FDR 5% (1,991 at FDR 1%, 698 at Bonferroni level; 3,140 with stratum-specific calibration). Because part of that excess reflects shared trait axes, we treat these counts as a more liberal sensitivity analysis under the separable null and keep the conservative *N* (0, 1) reference for discovery, with the class null (below) for interpreting individual hits.

#### Power of the screen for a single planted pair

To see what the screen can detect, we planted genetic-effect correlations between distal pairs of real enriched tags inside replicates of the simulated null (Methods; 12 replicates of 60 pairs, 720 pairs in four bins of LRCQ enrichment, *ρ* ∈ {0, 0.3, 0.5, 0.7, 0.9}, Table S8). Each planted tag keeps its realised signal, so the attainable correlation is capped by the tag’s own share of its projected genetic variance (realised *ρ* median 0.70 and 0.85 at nominal 0.7 and 0.9). No planted pair reached FDR 5%, with either the normal or the calibrated reference, while background false discoveries averaged 0.08 per screen. The median ∣*z*∣ rose only from 0.62 at *ρ* = 0 to 1.04 at *ρ* = 0.5, 1.54 at *ρ* = 0.7 and 1.84 at *ρ* = 0.9. At an empirical *P* < 10^−3^ against each replicate’s own null pairs, LRCP detected 15% of pairs with *ρ* ≥ 0.5 (from 5% in the least enriched bin to 20-21% in the two most enriched), and the naive correlation, size-adjusted in the same way, 12% (1-23%). This is the resolution floor of the targeted analysis seen genome-wide: single-pair SEs of about 0.3-0.5 give ∣*z*∣ ≈ 1-2 for *ρ* = 0.5-0.9, whereas 1.61 million tests need ∣*z*∣ above about 5.5. The 386 links with ∣*z*∣ ≥ 4.42 are therefore far stronger than a single separable correlation of this size can produce. They come from loci whose realised trait profiles are nearly collinear (∣*^ρ*∣ ≈ 1) along a shared trait axis, which is why we interpret narrow-class modules descriptively (below) and use gene-level aggregation to localise the signal.

#### Hits

Under the conservative normal reference, 437 pairs among 150 tags pass FDR 5% (216 at FDR 1%, 75 at Bonferroni level). All 437 keep *P* ≤ 1/2001, the floor of 2,000 draws, under the conditional parametric bootstrap in both orientations. Linking the 386 pairs with *P* < 10^−5^ gives 14 modules of 2-31 loci (Fig. 4c, Table 3). Each maps to a known biological process:

- **Platelet production** (31 loci): *ARHGEF3*, *GP6*, *F2R*, *DOCK8*, *TPM4*, *CD9*, *SIRPA* and others, acting on platelet volume and count in opposite directions.
- **Red-cell production** (24 loci): *HBS1L*, *CCND3*, *KIT*, *TFRC*, *CD164*, *TERT*, *IKZF1* and others, acting on red-cell volume.
- **Lipoprotein metabolism** (23 loci): *PCSK9*, *LDLR*, *APOB*, *ABCG8*, *CELSR2*, *APOC1*, *BCL3*, *CETP*, *LPL*, *LIPG*, *ABCA1*, joining LDL and HDL loci through the apolipoproteins.
- **Myeloid differentiation** (12 loci): *IRF8*, *NLRP12*, *ITGA4*, *BCL2A1*, *IKZF1*, *GSDMC*, acting on the monocyte versus neutrophil balance.
- **Bone mineral density** (8 loci): *WNT16*, *FZD7*, *CPED1*, *FMN2*.
- **Reticulocyte maturation** (7 loci): *HK1*, *SLFN14*, *TRIM58*, *SMOX*, *FREM3*, *RHOBTB3*.
- **Pigmentation** (7 loci): *HERC2*, *SLC45A2*, *SLC24A4*, *BNC2*, *DEF8*.
- **Alkaline phosphatase** (7 loci): *ALPL*, *GPLD1*, *ABCB11*, *RAP1GAP*.
- **Red-cell membrane** (6 loci): *ANK1*, *ARHGAP42*, acting on mean sphered cell volume.
- **Serum globulins** (4 loci): *TNFRSF13B*, *FCGR2A*, *FCGR2B*, *FCRLB*.
- **IL-6 signalling** (4 loci): *IL6R*, *CRP*, *LEPR*.
- **Kidney function markers** (3 loci): *SHROOM3*, *SHH*, *EPB41L5*, acting on cystatin C.
- Two smaller modules: *CBL*, *ABCC4*, *AK3* (platelet crit), and *COBLL1* × *KLF14* (triglycerides and HDL cholesterol).

**Table 3.** Genome-wide LRCP modules. Connected components of the 386 distal tag pairs with *P* < 10^−5^ among the 1,799 enriched tags. Genes are the nearest genes of the module’s loci (a gene can have more than one locus). Narrow trait class: at least one locus belongs to a trait-profile class with fewer than five effective traits; such modules are interpreted descriptively (Results). Signs refer to the coded allele of a reference locus, with the other loci aligned to it (Table S5). The full gene lists are in Supplementary Table S5.

| Module | Loci | Links | Genes (nearest gene of each locus) | Narrow trait class | Leading traits (sign) |
|---|---|---|---|---|---|
| 1 | 31 | 167 | *ARHGEF3*, *GP6*, *F2R*, *DOCK8*, *TPM4*, *CD9*, *SIRPA*, *CD226*, *GCSAML*, *BAZ2A*, *PSMD13*, *WDR66*, *MARK4*, *KALRN* and 14 others | yes | Mean platelet volume (-); Platelet count (+); Platelet distribution width (-) |
| 2 | 24 | 29 | *HBS1L*, *CCND3*, *KIT*, *TFRC*, *CD164*, *TERT*, *IKZF1*, *CCND2*, *NPRL3*, *PTPRC*, *RBM38*, *QKI*, *ATR* and 11 others | yes | Mean corpuscular volume (+); Mean corpuscular haemoglobin (+); Mean reticulocyte volume (+) |
| 3 | 23 | 79 | *PCSK9*, *LDLR*, *APOB*, *ABCG8*, *CELSR2*, *APOC1*, *BCL3*, *CEACAM19*, *BCAM*, *IRF2BP2*, *CETP*, *LPL*, *LIPG*, *ABCA1*, *ALDH1A2*, *ZNF259* | yes | Apolipoprotein A (-); Apolipoprotein B (+); LDL direct, adjusted by medication (+) |
| 4 | 12 | 44 | *IRF8*, *NLRP12*, *ITGA4*, *BCL2A1*, *IKZF1*, *GSDMC*, *CYP8B1*, *LPAR1*, *ST20-MTHFS*, *OR2K2* | yes | Monocyte percentage (+); Monocyte count (+); Neutrophil percentage (-) |
| 5 | 8 | 19 | *WNT16*, *FZD7*, *CPED1*, *FMN2*, *PTS*, *TMEM135*, *SEPT5*, *C2orf73* | yes | Heel bone mineral density (-); Heel quantitative ultrasound index (-); Heel BMD T-score (-) |
| 6 | 7 | 9 | *HK1*, *SLFN14*, *TRIM58*, *SMOX*, *FREM3*, *PELO*, *RHOBTB3* | yes | Reticulocyte percentage (+); Reticulocyte count (+); High light scatter reticulocyte percentage (+) |
| 7 | 7 | 11 | *HERC2*, *SLC45A2*, *SLC24A4*, *BNC2*, *DEF8*, *TCF25*, *SPATA33* | yes | Skin colour (-); Hair colour (natural, before greying) (-) |
| 8 | 7 | 8 | *ALPL*, *GPLD1*, *ABCB11*, *RAP1GAP*, *NBPF3* | no | Alkaline phosphatase (-); Phosphate (+); C-reactive protein (+) |
| 9 | 6 | 7 | *ANK1*, *ARHGAP42*, *OR10Z1*, *C9orf47* | yes | Mean sphered cell volume (-); Mean reticulocyte volume (-); Glycated haemoglobin (-) |
| 10 | 4 | 3 | *TNFRSF13B*, *FCGR2A*, *FCGR2B*, *FCRLB* | no | Non-albumin protein (+); Albumin/globulin ratio (-); Total protein (+) |
| 11 | 4 | 5 | *IL6R*, *CRP*, *LEPR* | no | C-reactive protein (-); Alkaline phosphatase (-); Albumin/globulin ratio (+) |
| 12 | 3 | 2 | *CBL*, *ABCC4*, *AK3* | no | Platelet crit (+); Platelet count (+); Mean sphered cell volume (-) |
| 13 | 3 | 2 | *SHROOM3*, *SHH*, *EPB41L5* | no | Cystatin C (+); eGFR, cystatin C (-); eGFR, creatinine + cystatin C (-) |
| 14 | 2 | 1 | *COBLL1*, *KLF14* | yes | Triglycerides (-); HDL cholesterol (+); Apolipoprotein A (+) |

The strongest pairs join loci that act in the same pathway: *ALPL* × *GPLD1* (*z* = −8.2, *^ρ* = −1.12), *IRF8* × *NLRP12* (*z* = −8.1), *SLC45A2* × *HERC2* (*z* = 7.9), *PCSK9* × *APOB* (*z* = −7.7) and *PCSK9* × *LDLR* (*z* = 7.3, *^ρ* = 0.99). Signs depend on the coded alleles, and they are biologically coherent: for all 62 hit pairs in the lipoprotein module the sign of *^ρ* equals the sign of the product of the two loci’s LDL-cholesterol Z scores, so LDL-raising alleles align.

#### Within-domain modules are reported descriptively

Most modules act on one trait domain (blood-cell indices, lipids, bone density, pigmentation), the situation in which the separable null is anti-conservative (simulation S7a). We therefore estimated profile classes over all 1,799 tags and, for each of the 437 hits, the class null with the pair’s own windows held out (§S8.5). Of the 437 hits, 405 have at least one locus in a class with fewer than five effective traits (*q*_eff_ 1.5-2.6). For the 366 pairs with both loci in such a class the class-null test has no power (∣*z*∣ ≤ 1.5), and their *^ρ* ≈ ±1 is read as shared membership of one trait axis, not as tested correlation; 13 of the 39 pairs with one such locus have class-null *P* < 0.05. For the other 32 pairs neither tag belongs to a profile class (fewer than five class members), so the class null falls back to the genome-wide trait covariance; their evidence rests on the separable null (*P* from 1.8 × 10^−16^ to 1.1 × 10^−5^, 7 below 5 × 10^−8^), and all have class-null *P* ≤ 5.4 × 10^−4^. They make up the alkaline phosphatase, serum globulin, IL-6, kidney and *CBL* modules, together with pairs in the red-cell, lipoprotein and pigmentation modules whose tags lie outside a profile class (for example *TFRC* × *CD164* and *LPL* × *ZNF259*).

#### Comparison with naive correlation

For the hit pairs the naive correlation is attenuated (median ∣*r*∣ 0.72 against ∣*^ρ*∣ 0.89; 5% of hits have ∣*^ρ*∣ > 1, a ratio-estimator effect, Methods), and naive correlation cannot separate them from the 1.14 million pairs it flags at FDR 5%.

#### Pathway validation

We mapped each tag to its nearest gene and asked whether linked gene pairs share a GO biological process of size 10-500 (Methods).

- 23.8% of the 386 LRCP links at *P* < 10^−5^ are co-annotated. Random distal pairs of the same enriched tags give 12.3% (1.9-fold; permutation *P* = 0.002). Rewiring the links while keeping each gene’s number of links gives 14.0% (*P* = 0.002).
- The 386 distal pairs with the largest naive ∣Z correlation∣ among the same tags are co-annotated less often (16.3%).
- At *P* < 10^−4^ (784 links) the enrichment is 1.7-fold (21.1% against 12.2%; rewired 13.0%).

LRCP links therefore point to shared biology beyond what naive phenome-wide similarity captures. Because nearest-gene mapping is approximate and links share hub loci, these P-values are indicative.

#### Gene-level burden tests within modules

We next asked which gene pairs carry each module’s signal (Methods, *Gene-level analysis*). A gene unit is a module gene in one LD window together with every enriched tag in that window whose nearest gene it is (124 units of 1-4 tags, median 1). Tags are oriented to the module member by the sign of their deconvolved trait-profile correlation. This step uses only the gene’s own window, never the partner window. For each distal window pair in a module, *^C_GH_* was tested against the conditional null given **Z***_B_*, using the local-moment *G_A_* and 2,000 parametric bootstrap draws in both orientations, with max-T family-wise error within the window pair (Table S5). This gave 899 gene pairs in 759 window pairs across the 14 modules. Of these, 298 contain a directly linked tag pair (*P* < 10^−5^); the other 601 are transitive, joined only through other module members.

- *Counts:* 757 gene pairs pass BH at 5% (bootstrap *P* at its floor of 1/2001 for 533 of them), 293 of 298 linked and 464 of 601 transitive; 762 pass max-T at 5% within their window pair. Because the modules and allele orientations were chosen from the same data, these counts are exploratory localisation within the modules, not confirmatory tests; confirmation needs modules and orientations frozen before testing in an independent panel. In the five modules without a narrow trait class, all 18 gene pairs pass BH (15 linked).
- *Aggregation:* for single-tag genes the burden reproduces the tag-pair statistic (317 pairs, *r* = 0.985 between *z* values). When a gene has several tags, aggregation strengthens the signal when they agree, for example *CELSR2* × *APOB* (tags per gene in parentheses: 1 and 3) ∣*z*∣ = 7.6 against 5.3 for the best tag pair, *CRP* (3) × *LEPR* (1) 7.0 against 5.2, *CETP* (4) × *LPL* (3) 6.2 against 4.9 and *SMOX* (2) × *RHOBTB3* 7.5 against 5.9. Across the 150 linked multi-tag pairs, burden ∣*z*∣ exceeds the best tag pair in 84 (median ratio 1.02).
- *Transitive pairs:* significant transitive pairs had a best tag pair below the linking threshold (median *P* = 7 × 10^−5^, range 10^−5^ to 10^−3^), for example *GCSAML* × *ANKMY1* and *BAZ2A* × *GP6* in the platelet module (both *z* = −5.2). The burden test thus fills in edges that the tag-level threshold missed, rather than creating new ones.

These tests are conditional on module membership, which was derived from the same summary statistics, so they localise each module’s signal to gene pairs and are not independent confirmation. We test the tag-level hits in independent cohorts in the next section.

### Replication in FinnGen and BioBank Japan

All 444 discovery GWAS come from the same UK Biobank participants, so we tested the 437 hits in two cohorts with no UK Biobank participants: FinnGen release 12 (57 traits matched to Pan-UKB traits; 35 clinical laboratory measurements, height, body-mass index, weight and 19 disease endpoints) and BioBank Japan (39 traits; 33 quantitative traits and 6 diseases). In each cohort we reran the whole procedure on the matched traits, with the cohort’s own LDSC plug-ins, the same 1,799 candidate tags and an ancestry-matched LD reference (1000 Genomes EUR for FinnGen and EAS for BioBank Japan; Methods). Both panels are much smaller than the discovery panel and consist mostly of blood biochemistry and blood-cell traits, so complete replication is not expected. As a ceiling, the same procedure on Pan-UKB restricted to the matched traits, which uses the discovery participants and is not a replication, gives two-sided *P* < 0.05 with the discovery sign for 389 (FinnGen traits) and 380 (BioBank Japan traits) of the 437 pairs.

|  | **FinnGen R12** | **BioBank Japan** |
|---|---|---|
| Matched traits (median *n*) | 57 (376,471) | 39 (145,502) |
| Testable hit pairs | 404 | 221 |
| Same sign as discovery | 365 (90%) | 168 (76%) |
| Two-sided *P* < 0.05, same sign | 229 | 69 |
| One-sided BH 5% | 237 | 57 |
| One-sided Bonferroni 5% | 28 | 5 |
| Correlation of *z* with discovery | 0.81 | 0.59 |
| Median-based λ, all distal pairs | 0.88 | 0.82 |

In FinnGen, 365 of the 404 testable pairs kept the discovery sign (binomial *P* = 9 × 10^−68^) and 237 passed a one-sided Benjamini-Hochberg threshold of 5% in the discovery direction (Fig. S5a, Table S10). The genomic-control factor over all 1.55 million distal pairs of the replication screen was 0.88, so the replication statistics are not inflated. The strongest discovery pair, *ALPL* × *GPLD1*, replicated with *z* = −5.0 (*^ρ* = −0.95), as did pairs in the lipoprotein (*PCSK9* × *LDLR*, *z* = 4.4) and myeloid modules (*ITGA4* × *IRF8*, *z* = −4.7). Pairs whose shared trait axis is missing from the replication panel could not replicate: no pigmentation trait was available, and *SLC45A2* × *HERC2* had *z* = 0.2. In BioBank Japan, 262 of the 1,799 tags were missing or had allele frequency below 1% in East Asians (for example the *PCSK9* tag), leaving 221 testable pairs. Of these, 168 kept their sign (*P* = 2 × 10^−15^) and 57 passed the one-sided BH threshold (Fig. S5b). Replication across ancestries was weaker, as expected from a smaller sample, fewer traits and different LD and allele frequencies, but it was well above chance. The genome-wide hits are therefore not an artefact of sample overlap within UK Biobank, and most of those whose traits were measured recur in an independent European cohort.

### Robustness

#### Trait set

We repeated the genome-wide screen on the 150 traits that remain after pruning to one trait per correlated cluster. This gives 121 pairs at FDR 5% (23 at Bonferroni level; median *λ* 0.91). Of the 437 main-analysis hits, 206 keep *P* < 10^−3^ and 61 pass FDR 5%. For the 476 pairs with *P* < 10^−3^ in both analyses, the signs agree in every case and the correlation of *^ρ* between the analyses is 0.995. Pruning removes most of the redundant blood-cell indices, so pigmentation loci (*SLC24A4*, *HERC2*, *SLC45A2*, *DEF8*) move to the top of the list. The module ranking therefore depends on the trait panel, as expected for an estimand defined over a trait population; the direction and size of *^ρ* do not.

#### Prescreen threshold

Restricting candidates to the 1,074 tags with LRCQ *q* < 0.01 (573,671 distal pairs) changes which tags are conditioned on in each window but leaves the statistics nearly unchanged. The screen finds 452 pairs at FDR 5% (71 at Bonferroni level; *λ*_median_ 0.89). Over the 930 pairs with *P* < 10^−3^ in both screens, *z* correlates at 0.998 and every sign agrees. Of the 284 main-analysis hits whose tags both pass the stricter threshold, 281 remain significant at FDR 5%. The prescreen only chooses which loci are tested and is not part of the conditional test, whose null holds for a candidate set chosen from LRCQ (simulation S4: prescreening on estimated enrichment keeps type-I error at or below the nominal level).

#### LD reference

We rebuilt LD for the 50 windows that hold the top 100 pairs from UK Biobank in-sample LD [Weissbrod 2020] and reran the confirmation. Every sign agrees with the 1000 Genomes EUR analysis, *z* correlates at 0.995, ∣*z*∣ is unchanged in the median (ratio 1.00; minimum ∣*z*∣ 3.5), and all 100 pairs keep bootstrap *P* ≤ 1/1001.

#### Sample-size convention

Using the effective sample size *n*_eff_ = 4/(1/*n*_cases_ + 1/*n*_controls_) instead of the total sample size for the 226 binary traits understates their *s_a_* = *n_a_h*^2^/*M*, because Pan-UKB estimated *h*^2^ with the total sample size (Methods). The screen is insensitive to this choice: *z* correlates at 0.99998 between the two conventions, and all 396 hits with *n*_eff_ are also main-analysis hits; the other 41 main hits have *q* between 0.050 and 0.060 with *n*_eff_ .

#### Other checks

No candidate tag lies in the MHC (chr6:25-34 Mb), so the MHC plays no part in the hits. All 444 GWAS are both-sexes analyses. For TP53 × MDM2, dropping SHBG changes nothing (above).

#### A 30-trait panel

With the 30 pruned traits of highest heritability *z*, the screen finds no pair at FDR 5% (*λ*_median_ 1.13). Three of the 437 main hits reach *P* < 10^−3^. This matches the power analysis (§S9): a 30-trait panel has too few genetically distinct traits, and it misses the blood-cell and lipid domains in which most hits act. LRCP needs phenome-scale panels.

#### Enrichment rectification

The genome-wide analysis does not depend on how *^w* is rectified. Candidates are chosen by LRCQ’s q-value from the unrectified OLS fit. The conditional test uses the local-moment trait covariance, and *^ρ* uses **R**-projected local variances, so *^w* enters neither the statistic nor its variance.

## Discussion

We defined the phenome-wide genetic-effect correlation, the row-wise counterpart of genetic correlation, and developed LRCP to estimate it from GWAS summary statistics. Three of our results bear on how such correlations should be studied.

### Naive phenome-wide correlations are not evidence on their own

Biobank GWAS share samples and genetically correlated traits, so the Z scores of one SNP across hundreds of traits carry far less information than their number suggests. In Pan-UK Biobank, 452 GWAS behave like 58 independent ones, and the more pleiotropic a SNP is, the fewer effective traits its correlations rest on. Naive tests are therefore least reliable for the highly pleiotropic variants that attract the most interest, and correlations of unsigned statistics such as — log_10_ *P* are biased upwards by differences in power between traits. These results apply to any analysis that clusters or correlates variant-by-trait association matrices, including pleiotropy-profile clustering [Udler 2018; Smith 2024], DeGAs [Tanigawa 2019] and PDR [Ballard & O’Connor 2022]. The correction is inexpensive: the effective number of traits *q*_eff_ = tr **M***_k_* tr **M***_l_*/ tr(**M***_k_***M***_l_*) is computable from LDSC output, and an empirical null from random distal pairs is easy to build.

### LRCP separates genetic-effect correlation from LD and GWAS dependence

Writing the expected Z-score cross-product as the sum of a sampling-noise, a tagging and a correlation component makes the three sources of a statistical link explicit. For distal pairs the estimator collapses to an LD deconvolution of the trait-averaged cross-product matrix, which is fast, needs only the candidates’ Z scores and LD, and has exact standard errors under any trait dependence. In simulations it removes the LD attenuation of naive correlations and holds the type-I error at its nominal level where naive tests reject half the null pairs.

### Resolution is set by genetically distinct traits

LRCP’s precision for a single pair has a floor of about 1/ *q_G_*, the number of genetically distinct traits, no matter how strongly enriched the SNPs are. With current Pan-UK Biobank data, correlations below about 0.3 are below resolution for a single pair (Supplementary Note §S9.4), and the TP53–MDM2 estimate (*^ρ* = 0.90, SE 0.59; bootstrap P = 0.015, but P = 0.18 under the class null, whose classes span fewer than five effective traits) illustrates the limit. Discovery is better done on aggregates, such as gene and block pairs or clusters of enriched clumps, where coherent correlations add up, and power grows with trait diversity: adding GWAS of new trait domains (molecular traits, imaging, diseases from other cohorts) raises *q_G_* far more than adding further correlated traits from one domain.

Simple tests of whether a SNP’s Z scores vary more across traits than expected, such as a Tukey range test or a sample-variance test over traits, assume independent traits. Under trait correlation and sample overlap, the effective number of genetically distinct traits in Pan-UKB is about 10-11, not 444, so they are invalid (Supplementary Note §S4.5). The conditional and max-T tests replace them.

### Biological interpretation

Genome-wide, the loci LRCP links fall into modules that each correspond to a known process: platelet and red-cell production, lipoprotein metabolism, lineage choice in myeloid cells, bone mineralisation, pigmentation, release of alkaline phosphatase and IL-6 signalling. That the method recovers such modules from summary statistics alone, without gene annotation, is its main practical use. Most modules act on one trait domain, and there LRCP shows shared membership of a trait axis rather than a tested correlation (§S8.5). Linked gene pairs share GO biological processes 1.9 times more often than random pairs of enriched loci. They also share them more often than the most similar pairs by naive correlation. The TP53 and MDM2 variants affect the same traits in the same direction, mainly blood pressure, red-cell indices and body composition, consistent with their roles in one feedback loop. The rare TP53 variant is known to reduce *TP53* mRNA processing [Stacey 2011], and the MDM2 variant is in the gene whose product degrades p53. A shared effect on p53 activity is the simplest explanation, but LRCP shows similarity of effects, not mechanism, and a functional study is needed.

### Limitations

#### Separable effects

The model assumes the SNP-by-trait effect covariance factorises into SNP and trait parts. When two loci act only on one narrow trait domain, their effective number of traits is tiny, and in simulation (S7a) tests against *ρ* = 0 reject 9-31% of null pairs instead of 5%. Correlations among such loci, including the lipid controls, are best read as within-domain similarity over the observed traits. A class null built from held-out loci of the same trait-profile class (§S8.5) addresses this directly, but has no power when the class spans fewer than about five effective traits.

#### Plug-ins

LRCP inherits errors in LDSC intercepts, genetic covariances and LRCQ enrichment. In simulation, 20% errors in *h*^2^ bias *^ρ* by about 6% and intercept errors matter little, but ignoring genetic correlation between traits inflates type-I error up to 32%, so genome-wide cross-trait LDSC is required.

#### Candidate set

Which pairs are found depends on which loci the LRCQ prescreen admits: loci it misses are never tested, and a stricter prescreen trades discovery for fewer tests (Robustness).

#### LD reference

Deconvolution among candidates in partial LD is sensitive to the LD reference. The targeted analysis used UK Biobank in-sample LD; the genome-wide screen used 1000 Genomes EUR and therefore relies on the conditional test. In simulation (S5), a 1000 Genomes EUR reference doubled the type-I error of the GLS test, while the conditional test stayed calibrated, and the top genome-wide pairs gave the same results with UK Biobank in-sample LD (Robustness). Out-of-sample references should be used with clumping to *r*^2^ < 0.5 and the **R**-projected denominators.

#### Bootstrap calibration

The conditional bootstrap is calibrated at the TP53 × MDM2 configuration (3.3-4.5% at 5%), but in a numerical check of the gene-burden null (Supplementary Note, Appendix A) it reached 7.3% at suggestive. *q* = 300. Single-pair bootstrap P-values near 0.01 should therefore be read as

#### Genome-wide calibration

A per-pair bootstrap is infeasible for 1.6 million pairs, so the screen uses the *N* (0, 1) reference, which a simulated null of the whole screen shows to be conservative. Against that simulated null the bulk of observed statistics departs in both directions (*λ*_cal_ 1.21 overall, from 1.65 to 0.67 across deciles of genetic share), and the extreme tail sits mostly in pairs whose variance is mostly genetic. We attribute this to loci concentrating on shared trait axes, which the separable null does not model, so the 3,154 pairs at calibrated FDR 5% are a liberal sensitivity count and the 437 pairs under the normal reference our primary result. Under the Benjamini-Yekutieli procedure, valid under any dependence among the pair statistics, 126 of the 437 remain. A null that models trait-axis structure genome-wide, for example a class null at scale, would sharpen this.

#### Rare variants

The enrichment of rare variants such as rs78378222 is estimated imprecisely, which carries into *^ρ*.

#### Local pairs

We focused on distal pairs. Within an LD block LRCP must also separate *ρ* from the LD and tagging components, which needs the full regression or likelihood and is less precise.

#### Assortative mating

Assortative mating induces weak long-range LD between loci that affect the mated-on traits (height, education). For distal pairs this produces a correlation of Z scores that LRCP, given an LD reference without that structure, attributes to *ρ*. Modules acting on such traits should be checked with within-family GWAS.

#### Population

All analyses are in European-ancestry data.

### Outlook

Beyond discovery, *^ρ* could be used in three ways. The most direct is as a joint prior for multi-trait fine-mapping and genomic prediction. A multivariate ridge regression that fits all traits at once with the penalty **K***_g_* ⊗ **K***_s_*, *^β*(**K**) = (**I***_q_* ⊗ **X**^⊤^**X** + **K***_g_* ⊗ **K***_s_*)^−1^(**I***_q_* ⊗ **X**^⊤^)**y**, is the posterior mean under the prior 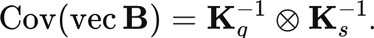. The trait side **K**_g_^−1^ holds per-causal-variant heritabilities and genetic covariances, 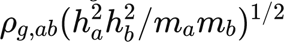. The SNP side **K**_g_^−1^ has the enrichments *w_i_* on its diagonal (a unit diagonal when enrichment is uniform) and 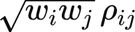 off the diagonal. This is the matrix-normal model of LRCP (Methods), so LRCQ’s *^w* and LRCP’s **^R** β can supply the SNP-side covariance, which such methods otherwise set to the identity or give a vague prior. The other two uses are as a pairwise functional-similarity score and as input to matrix factorisation of the variant-by-trait matrix (DeGAs [Tanigawa 2019], FactorGO [Zhang 2023]). LRCP and LRCQ together give a two-step map from phenome-wide summary statistics to functional relationships: LRCQ finds the variants with outsized phenome-wide effects, and LRCP asks which of them act alike. As biobanks add diverse trait domains and larger samples, these correlations will become resolvable for individual pairs; until then gene- and module-level aggregation is the practical route to discovery.

## Methods

### Genetic model

We consider *q* traits measured on standardised scales, with *m* standardised SNP genotypes. For a cohort of *n*individuals,

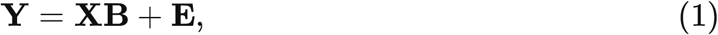

where **B** (*m* × *q*) holds the joint (causal) per-standard-deviation effects. We treat **B** as random with E[**B**] = 0 and two second-moment structures: the SNP-side covariance **Σ***_β_* = E[**BB**^⊤^]/*q* (*m* × *m*) and the trait-side covariance **Σ***_g_* = E[**B**^⊤^**B**]/*m* (*q* × *q*). The diagonal of *m***Σ***_g_*contains the SNP heritabilities *h*^2^ and its off-diagonal entries the genetic covariances *h_ab_*, which LDSC [Bulik-Sullivan 2015a,b] and HDL [Ning 2020] estimate. For simulation and likelihood-based inference we further assume that **B** is matrix normal [Dawid 1981] with these two covariance structures; its explicit form is given after eq. (2). The moment results below need only the first two moments (Supplementary Note §S1.2).

We decompose **Σ***_β_* into an enrichment vector and a correlation matrix,

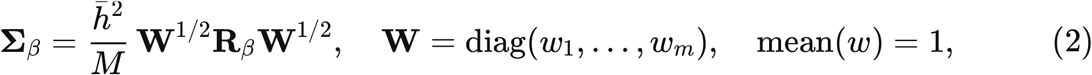

where *h*^ˉ2^ is the average heritability across traits and *M* the number of SNPs over which heritability is spread. *w_i_* is SNP *i*’s *phenome-wide heritability enrichment*, the target of the companion method LRCQ [Chen 2026]. The (*i*, *j*) entry *ρ_ij_*of **R***_β_* is the **phenome-wide genetic-effect correlation** of SNPs *i* and *j*: the correlation, across a population of traits, of the two SNPs’ joint effects. It is the transpose of the genetic correlation *r_g_*, which correlates two traits’ effects across SNPs. In matrix-normal form, **B** ∼ MN(0, **W**^1/2^**R***_β_***W**^1/2^, **H**/*M*) with **H** = (*h_ab_*), that is, Cov(vec **B**) = (**H**/*M*) ⊗ **W**^1/2^**R***_β_***W**^1/2^, so that the common scale *h*^ˉ2^/*M* of eq. (2) enters once. Biologically, *ρ_ij_* =f 0 arises when two variants act through shared intermediate molecular mediators. If mediators **m** = **Px** + **e** influence traits through **y** = **Qm** + **u**, then **B** = **P**^⊤^**Q**^⊤^ and **Σ***_β_* ∝ **P**^⊤^(E **Q**^⊤^**Q**)**P**, so SNPs that load on the same mediators (for example the same gene’s expression, or two genes in one pathway) have correlated rows of **B** (Supplementary Note §S2.3; Fig. 1a).

For a gene pair (*g*, *h*), we define the gene-level correlation as the enrichment-weighted mean of SNP-pair correlations,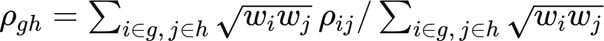.

### From joint effects to GWAS summary statistics

With LD matrix **R**, the GWAS Z scores of trait *a* satisfy 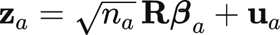 where **u***_a_* ∼ *N* (0, **R**) up to confounding. The **u***_a_* are correlated across traits through sample overlap and phenotypic correlation: Cov(**u***_a_*, **u***_b_*) = *c_ab_***R**, with *c_aa_* the LDSC intercept and *c_ab_* the cross-trait LDSC intercept [Bulik-Sullivan 2015b]. Theorem 1 of the Supplementary Note (§S1.4) shows that

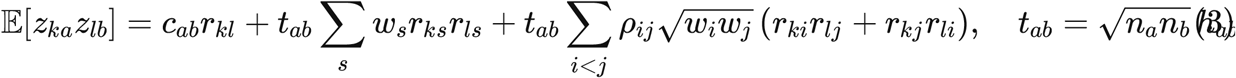

The three terms separate the sources of a “statistical link” between SNPs *k* and *l*: shared sampling noise through LD (first term), true effects tagged by both SNPs (second term), and genetic-effect correlation between causal SNPs tagged by *k* and by *l* (third term). Each unordered causal pair enters once; counting it twice halves the regression slope (Supplementary Note §S1.5; simulation S8). In matrix form, 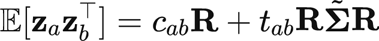, with 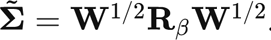.

### Distal pairs: LD-deconvolved cross-product estimator

Most of the biology we target involves SNPs in different LD blocks or on different chromosomes (as with TP53 and MDM2). Let *A* and *B* be two sets of SNPs with no LD between them (*r_kl_* = 0 for *k* ∈ *A*, *l* ∈ *B*). Then the first two terms of (3) vanish for every cross pair, and summing over traits gives

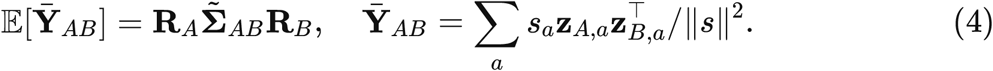

**Y**^ˉ^ is the trait-averaged cross-product matrix of_∼_Z scores, scale_∼_d to the per-unit genetic signal. Equation (4) shows that LD acts on both sides as a *blur* on **Σ**. Restricting **Σ***_AB_* to candidate causal SNPs *P_A_* ⊂ *A* and *P_B_* ⊂ *B* (below), the regression of vec(**Y**^ˉ^ *_AB_*) on (**R***_B_*_,⋅_*_PB_*⊗ **R***_A_*_,⋅_*_PA_*) under the working covariance Cov(vec **Y**^ˉ^) ∝ **R***_B_* ⊗ **R***_A_* (exact when the per-SNP signal is small relative to sampling noise) has the generalised least-squares solution

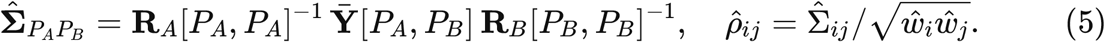

The estimator uses only the candidate SNPs’ Z scores and their LD submatrix and costs two small linear solves per window pair. (The derivation is in Supplementary Note §S4.6: **R**^−1^**R***_A_*_,⋅_*_P_* selects the identity columns.) When *P* is the whole window, (5) reduces to full LD deconvolution of **Z**, which is unstable in high LD. Prescreening on *^w* therefore serves both as biology (no variance implies no covariance) and as regularisation.

#### Tight LD

Per-SNP effects are not identifiable among SNPs in near-perfect LD: *D* = **R** ∘ **R** is numerically singular among HapMap3 SNPs. We therefore prune each candidate set greedily to tags with pairwise *r*^2^ < 0.5, keeping the highest-*^w* SNP, before inverting **R**[*P*, *P*]. Each tag *t* then estimates the covariance of its **R**-*projected* effect 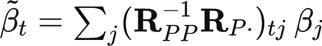, i.e. the tag plus its linked SNPs (Supplementary Note §S3.11, §S4.7). So that numerator and denominator refer to the same projected effects, the variances in the denominator come from the same projection: 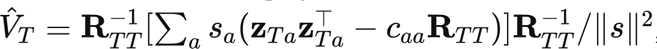, and *^p_tu_* = 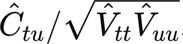 the level of LD clumps (“omni-loci”), using LRCQ’s tag-set enrichment.

#### Cross-trait products

Products *z_ka_z_lb_* with a ≠ *b* have expectation *t_ab_*(**RΣ**^∼^ **R**)*_kl_* and add information when traits are genetically correlated. The general estimator weights all trait pairs by *t_ab_*(Supplementary Note §S4.2). We use same-trait products (*a* = *b*) as the default because they do not depend on estimated *h_ab_*.

### Local pairs: full regression and likelihood

For SNPs within the same or adjacent blocks, all three terms of (3) are present. We regress Z cross-products within a sliding window of three LD blocks on the LD cross components {*r_kl_*, ∑*_s_ w_s_r_ks_r_ls_*, *r_ki_r_lj_* + *r_kj_r_li_*}, with *c_ab_*, *h_ab_* and *w* fixed at their stage 1-3 estimates. Estimates are kept only for SNPs in the middle block [Chen 2026]. We provide ordinary and weighted least squares, with weights from the Isserlis variance Var(*z_k_z_l_*) = *σ_kk_σ_ll_* + *σ*^2^ (Supplementary Note §S3), and a Gaussian maximum-likelihood estimator under the matrix-normal model (lrcp_mle). The MLE maximises the likelihood of the screened candidates’ Z scores with the stage 1-3 plug-ins fixed, and takes SEs from the numerical Hessian. The approach follows the eigen-decomposed likelihood of HDL [Ning 2020]. It costs minutes per candidate set even at *q* = 30, so we use it only on small screened sets. In simulations it is more precise than GLS, but its Hessian SEs are too small (95% coverage 63-70%), so tests use GLS and the conditional test (Results, S1-MLE).

### Four-stage procedure

1. **Heritability and co-heritability.** *^h_a_*^2^ and *^h_ab_* by univariate and cross-trait LDSC on genome-wide HapMap3 SNPs [Bulik-Sullivan 2015a,b] (HDL [Ning 2020] as a sensitivity analysis).
2. **Intercepts**. *^c_aa_* and *^c_ab_* from the same LDSC fits, held fixed genome-wide (per-window intercepts make *w* unidentifiable; Supplementary Note §S3).
3. **Enrichment** *^w*. Tag-set LRCQ by OLS, unrectified, clipped at 0 where a variance must be positive [Chen 2026].
4. **Correlation** *^ρ*. Candidate sets are LD-clump tags with evidence of enrichment, {*i* : *^w_i_* > 0} at LRCQ FDR *q* < 0.05 (prescreening simulation S4, Supplementary Table S4). The genome-wide set (1,799 tags) used the companion paper’s score-type LRCQ q-values. Distal pairs by (5), local pairs by the windowed regression or MLE.

Table 4 summarises the unknowns of each stage, how they are estimated and where the Pan-UKB application stands.

**Table 4.**
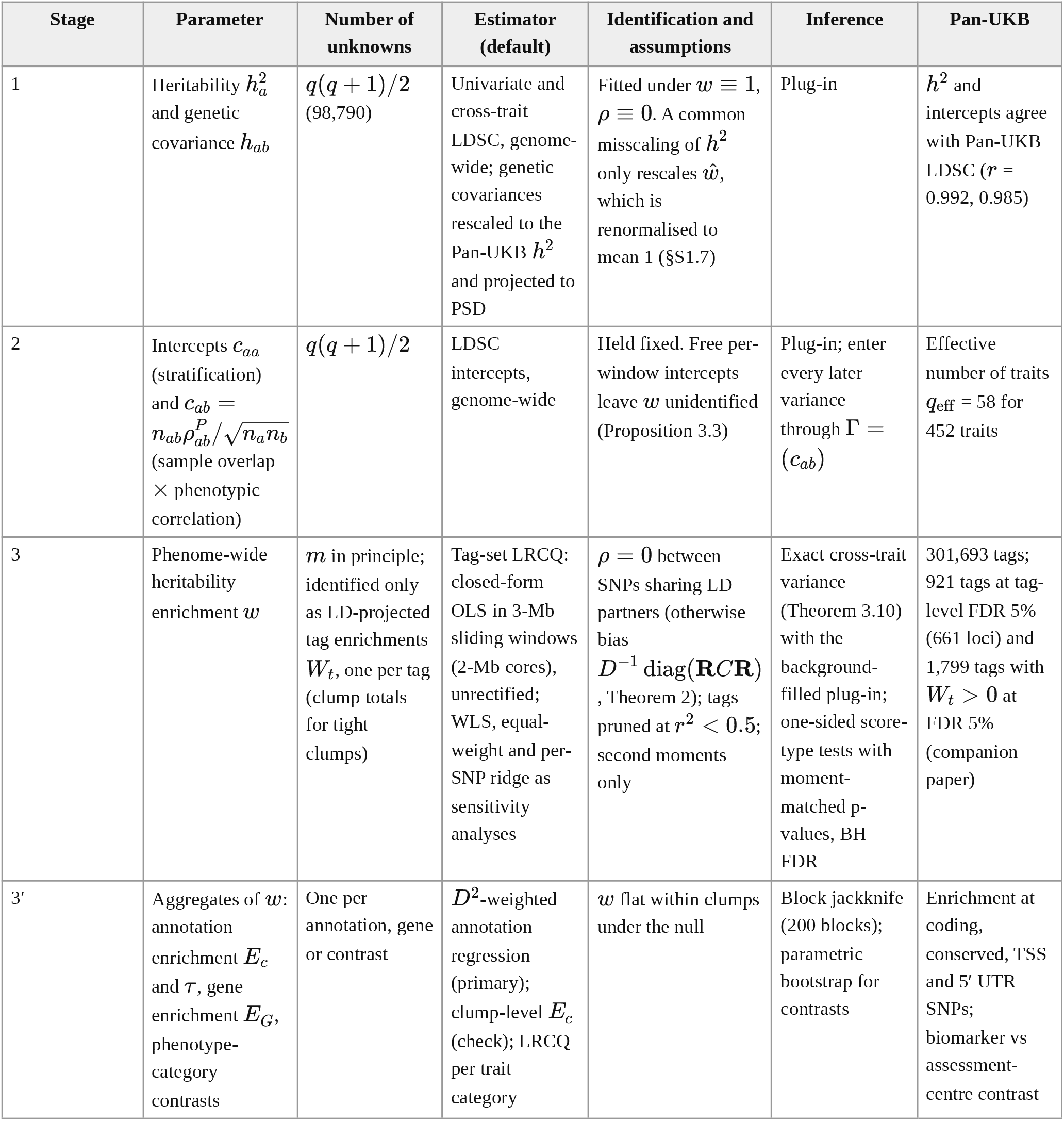
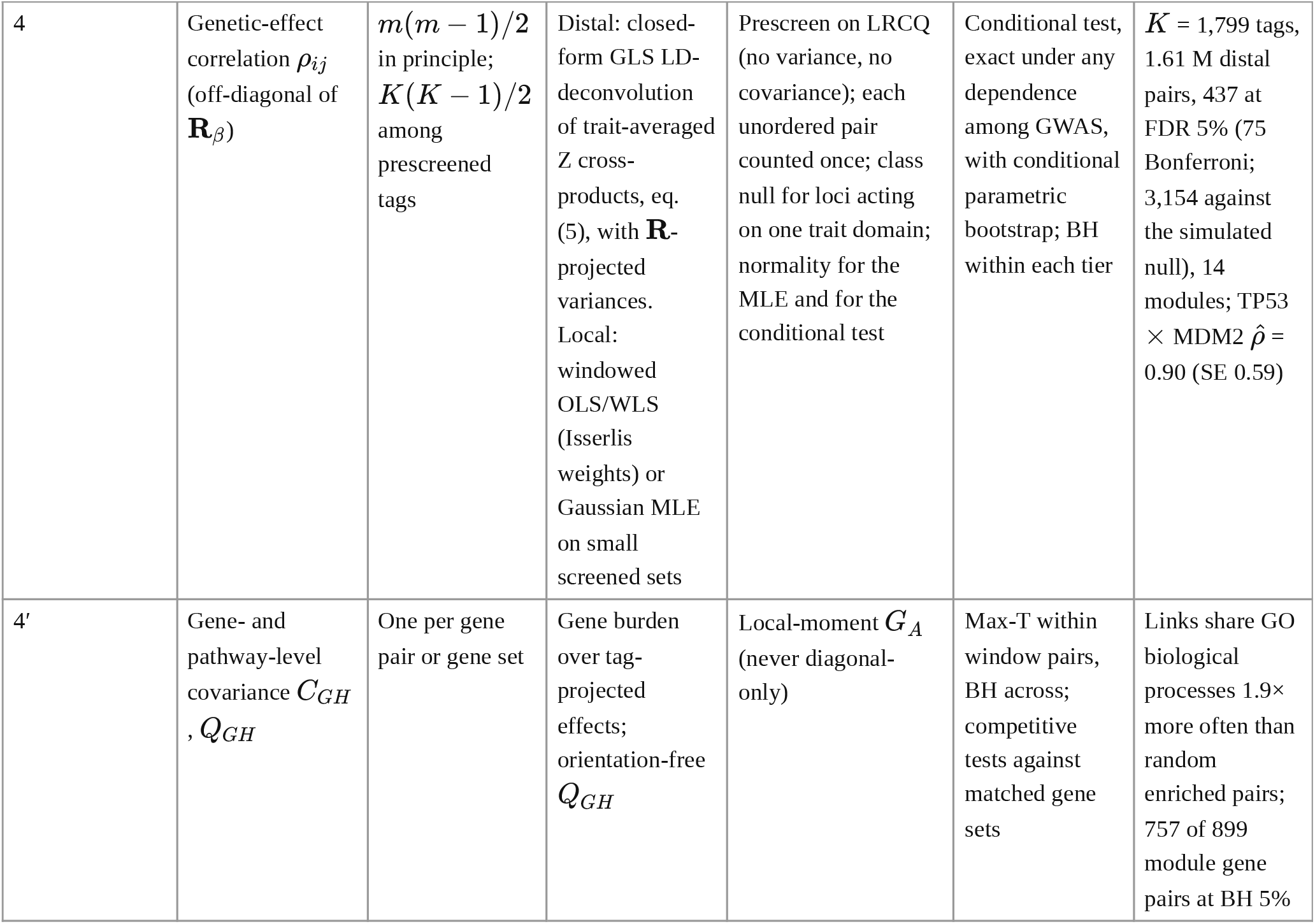
Unknown parameters by stage: estimator, identification, inference and Pan-UKB results (Methods, “Four-stage procedure”).

| Stage | Parameter | Number of unknowns | Estimator (default) | Identification and assumptions | Inference | Pan-UKB |
|---|---|---|---|---|---|---|
| 1 | Heritability h<sub>a</sub><sup>2</sup> and genetic covariance h<sub>ab</sub> | q(q + 1)/2 (98,790) | Univariate and cross-trait LDSC, genome-wide; genetic covariances rescaled to the Pan-UKB h<sup>2</sup> and projected to PSD | Fitted under w ≡ 1, ρ ≡ 0. A common misscaling of h<sup>2</sup> only rescales ŵ, which is renormalised to mean 1 (§S1.7) | Plug-in | h<sup>2</sup> and intercepts agree with Pan-UKB LDSC (r = 0.992, 0.985) |
| 2 | Intercepts c<sub>aa</sub> (stratification) and c<sub>ab</sub> = $n_{ab}\rho^{P}_{ab}/\sqrt{n_a n_b}$ (sample overlap × phenotypic correlation) | q(q + 1)/2 | LDSC intercepts, genome-wide | Held fixed. Free per-window intercepts leave w unidentified (Proposition 3.3) | Plug-in; enter every later variance through Γ = (c<sub>ab</sub>) | Effective number of traits q<sub>eff</sub> = 58 for 452 traits |
| 3 | Phenome-wide heritability enrichment w | m in principle; identified only as LD-projected tag enrichments W<sub>t</sub>, one per tag (clump totals for tight clumps) | Tag-set LRCQ: closed-form OLS in 3-Mb sliding windows (2-Mb cores), unrectified; WLS, equal-weight and per-SNP ridge as sensitivity analyses | ρ = 0 between SNPs sharing LD partners (otherwise bias $D^{-1}\,\mathrm{diag}(\mathbf{R}C\mathbf{R})$, Theorem 2); tags pruned at r<sup>2</sup> < 0.5; second moments only | Exact cross-trait variance (Theorem 3.10) with the background-filled plug-in; one-sided score-type tests with moment-matched p-values, BH FDR | 301,693 tags; 921 tags at tag-level FDR 5% (661 loci) and 1,799 tags with W<sub>t</sub> > 0 at FDR 5% (companion paper) |
| 3′ | Aggregates of w: annotation enrichment E<sub>c</sub> and τ, gene enrichment E<sub>G</sub>, phenotype-category contrasts | One per annotation, gene or contrast | D<sup>2</sup>-weighted annotation regression (primary); clump-level E<sub>c</sub> (check); LRCQ per trait category | w flat within clumps under the null | Block jackknife (200 blocks); parametric bootstrap for contrasts | Enrichment at coding, conserved, TSS and 5′ UTR SNPs; biomarker vs assessment-centre contrast |
| 4 | Genetic-effect correlation ρ<sub>ij</sub> (off-diagonal of $\mathbf{R}_{\beta}$) | m(m − 1)/2 in principle; K(K − 1)/2 among prescreened tags | Distal: closed-form GLS LD-deconvolution of trait-averaged Z cross-products, eq. (5), with $\mathbf{R}$-projected variances. Local: windowed OLS/WLS (Isserlis weights) or Gaussian MLE on small screened sets | Prescreen on LRCQ (no variance, no covariance); each unordered pair counted once; class null for loci acting on one trait domain; normality for the MLE and for the conditional test | Conditional test, exact under any dependence among GWAS, with conditional parametric bootstrap; BH within each tier | K = 1,799 tags, 1.61 M distal pairs, 437 at FDR 5% (75 Bonferroni; 3,154 against the simulated null), 14 modules; TP53 × MDM2 ρ̂ = 0.90 (SE 0.59) |
| 4′ | Gene- and pathway-level covariance C<sub>GH</sub>, Q<sub>GH</sub> | One per gene pair or gene set | Gene burden over tag-projected effects; orientation-free Q<sub>GH</sub> | Local-moment G<sub>A</sub> (never diagonal-only) | Max-T within window pairs, BH across; competitive tests against matched gene sets | Links share GO biological processes 1.9× more often than random enriched pairs; 757 of 899 module gene pairs at BH 5% |

Table 4. Unknown parameters of the LRCQ-LRCP model, how each is estimated, and status in Pan-UKB. Inputs are the Z-score matrix **Z** (*m* × *q*; *m* = 1,094,844 HapMap3 SNPs, *q* = 444 Pan-UKB EUR traits), total sample sizes *n_a_* (cases plus controls), *M* = 1,094,844 and LD **R** (1000 Genomes EUR in 1,383 3-Mb windows; UK Biobank in-sample LD as a sensitivity analysis). Each stage holds the previous stages’ estimates fixed

### Inference

The sampling unit for *ρ* is the *trait*, and traits are dependent. As a result, the null variance of any trait-summed statistic depends on the full *q* × *q* covariance **M***_k_* of each SNP’s Z vector: Cov(*z_ka_z_la_*, *z_kb_z_lb_*) = *M_k_*_,*ab*_*M_l_*_,*ab*_ under *ρ* = 0 (Isserlis). We showed by simulation that ignoring this inflates type-I error of naive tests to 20-67% under realistic UK Biobank trait structure (Supplementary Table S9 and Fig. 1f; effective number of traits *q*_eff_ = tr **M***_k_* tr **M***_l_*/ tr(**M***_k_***M***_l_*)). Standard errors use the exact Isserlis covariance of the cross-products with **^M** *_k_* built from stage 1-3 plug-ins (Supplementary Note, Theorem 3.10), with *^w* clipped at 0 so that **R** Diag(*^w*)**R** is positive semi-definite. GLS tests use the SE under *ρ* = 0 and confidence intervals the plug-in SE, which adds the second Isserlis term Cov(*z_ka_*, *z_lb_*) Cov(*z_la_*, *z_kb_*) evaluated at *^ρ* and is needed for ∣*ρ*∣ > 0.4. For pairs of loci that act on a narrow trait domain, the trait covariance in these variances is replaced by a class covariance *^V*_class_, estimated from other enriched loci of the same trait-profile class with the tested loci and their blocks held out (class null, Supplementary Note §S8.5), and *q*_eff_ (*^V*_class_) is reported with each pair. The conditional test of the gene-level section is the primary test, reported with the ratio *R_AB_* of its variance to the GLS variance. P values for the primary conditional test come from the conditional parametric bootstrap (both orientations, larger P; §S8.2a); GLS P values use *N* (0, 1). In the genome-wide screen, where a bootstrap per pair is infeasible for 1.61 million pairs, the conditional statistic is referred to *N* (0, 1), which is conservative for this self-normalised statistic, and calibrated against a parametric null of the whole screen (*Genome-wide design* below). Multiple testing is controlled by the Benjamini-Hochberg FDR within each analysis tier (targeted; genome-wide among prescreened SNPs).

### Power and resolution

Var *^C_ij_* = [1/*q_P_* + *s*(*w_i_* + *w_j_*)/*q_PG_* + *s*^2^(*w_i_w_j_* + *C* ^2^)/*q_G_*]/*s*^2^ (Supplementary Note §S9). It involves two effective trait counts: 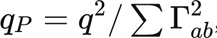, set by sample overlap and phenotypic correlation, which governs the null, and 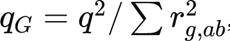, the genetic count, which governs power at large enrichment. A floor *r_b_* on the genetic correlation between trait domains caps *q_G_* near 1/*r*^2^, so adding distinct trait domains helps more than adding traits. Genome-wide we test all tags with LRCQ *q* < 0.05 (*K* = 1,799) and control FDR by BH, rather than prescreening to *K* ≤ 100 with Bonferroni (§S9.4). We report the detectable ∣*ρ*∣ at 80% power for each analysis and treat ∣*ρ*∣ < 0.3 as below the resolution of current Pan-UKB data. At *n* = 4 × 10^5^, *q*_eff_ = 300, *h*^2^ = 0.1 and *K* = 100, power at *ρ* = 0.5 is 0.83 for *w* = 50 but 0.18 for *w* = 20.

### Gene-level analysis

A gene *G* is represented by a weight vector *v_G_* over the enriched tags of one window (in the genome-wide analysis, the tags whose nearest protein-coding gene is *G*, with signs from the orientation step below). For genes *G* (window *A*) and *H* (window *B*), the gene-burden genetic-effect covariance *C_GH_* = *v*^⊤^*Cv_H_* over tag-projected effects is estimated by 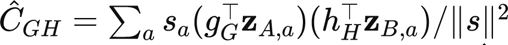, with *g_G_* = **R***_A_*[*T*, *T*]^−1^*v_G_* and *h_H_* = **R***_B_*[*T*, *T*]^−1^*v_H_* (Supplementary Note §S8). The sign of *^C_GH_* depends on the allele coding of each tag. We therefore orient the tags of a gene to a reference tag by the sign of the correlation between their deconvolved trait profiles; this uses only the gene’s own window, never the partner window, so it does not affect the null of the cross-window test. For block pairs (targeted analysis) we also report the orientation-free statistic 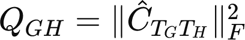, with a Liu-Satterthwaite reference.

#### Conditional test, exact under any dependence among GWAS

Conditional on **Z***_B_*, *^C* is a linear function of **Z***_A_*. Under *H*_0_ (no cross-window genetic-effect covariance) and Gaussian Z scores it is therefore exactly Gaussian with

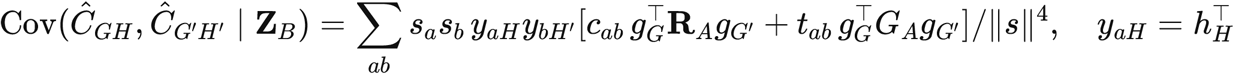

which needs only window *A*’s second moments (LD, intercepts, *h_ab_* and the local genetic covariance *G_A_*). *G_A_* is estimated by the local moment 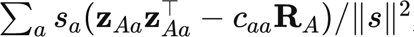, clipped to be positive semi-definite and shrunk towards **R** Diag(*^w*_+_)**R** when *q*_eff_ < 20. Because this shares **Z***_A_* with the numerator, P-values come from a conditional parametric bootstrap (redraw **Z***_A_* with **Z***_B_* fixed; both orientations, larger P reported; Supplementary Note §S8.2a). A diagonal-only *G_A_*, which assumes no correlation between the causal SNPs of a gene, must not be used for gene, pathway or max-T tests. Critical values for the maximum ∣*T* ∣ over all gene pairs in a window pair are simulated from this Gaussian, giving exact family-wise error within the window pair whatever the sample overlap and genetic correlation among traits. Window pairs are then combined by Benjamini-Hochberg [Benjamini & Hochberg 1995] (Benjamini-Yekutieli [Benjamini & Yekutieli 2001] as a conservative check).

#### Genome-wide design

1. *Screen.* Every distal pair of enriched tags is tested at tag level with the conditional statistic above, conditioning on the other enriched tags of each window and using the larger of its two orientation variances and the plug-in local moment of each window. (2) *Calibration.* Because the local moment is estimated from the same Z scores as the numerator, this statistic is self-normalised and light-tailed under *H*_0_, so its *N* (0, 1) reference is conservative. We calibrate it with a parametric null of the whole screen: every window is redrawn independently as 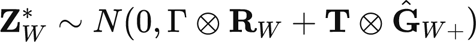, which keeps each window’s LD, trait covariance and local signal but has no genetic-effect covariance between windows, and the full screen is recomputed, including 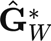. Pooling the null ∣*z*∣ over 200 replicates (322 million null pairs) gives the calibrated *P* = (1 + #{∣*z*^∗^∣ ≥ ∣*z*∣})/(*N* ^∗^ + 1). A stratified version, with strata defined by deciles of the pair’s share of genetic variance in its conditional variance (fixed from the observed data), checks that pooling across heterogeneous pairs does not matter. Discovery uses BH on the conservative *N* (0, 1) P-values. BH on the calibrated P-values is reported as a liberal sensitivity analysis, because the simulated null is separable and pairs of loci on a shared trait axis exceed it without a specific shared mechanism (§S8.5). (3) *Confirmation.* The top pairs are re-tested with the pair-specific conditional parametric bootstrap (both orientations, larger P) and with the class null. (4) *Modules.* Pairs below a P threshold form the edges of a graph whose connected components are the modules. (5) *Gene burden within modules.* Gene units are each module gene’s enriched tags in one window, oriented as above. *C^GH* is tested for every distal window pair of the module, with max-T within the window pair and BH across gene pairs. (6) *Power.* We planted genetic-effect correlations between distal pairs of real enriched tags in replicates of the null (spike-in): for a planted tag *i*, the share *c_i_* = 1/(**Σ**^−1^)*_ii_* of its projected genetic variance (**Σ** = **R**^−1^**G ^R**^−1^) is drawn as a separate factor 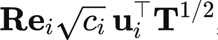, and the factors of the two tags of a pair are correlated so that their projected effects have correlation *ρ*. This keeps each tag’s realised signal and tests the planted pairs among all 1.61 million pairs of the replicate. Tags with *c_i_*/Σ*_ii_* ≥ 0.5 were eligible (1,792 of 1,799); each replicate planted 60 pairs in distinct windows, cycling over four bins of LRCQ enrichment (*W* in [0, 50), [50, 100), [100, 150), ≥ 150) and *ρ* ∈ {0, 0.3, 0.5, 0.7, 0.9}, with the planted correlation capped at 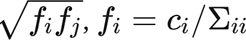 (12 replicates). 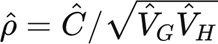 has a ratio bias of about 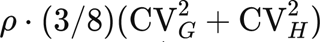 and can exceed 1 in absolute value when the variances are noisy. We therefore test and rank on *^C* and its *z*, and report *^ρ* descriptively.

### Comparators

i. Naive Pearson correlation of Z scores across traits with the Fisher test on *q* − 3 degrees of freedom. (ii) The same with *q*_eff_ degrees of freedom. (iii) Correlation of − log_10_ *P* (as in the motivating MAGMA analysis). (iv) In simulations, an oracle: the empirical correlation of the true rows of **B**.

### Simulations

Phenome-wide GWAS were simulated with lrcpq::simulate_lrcpq from **B** ∼ MN(0, **W**^1/2^ **R***_β_* **W**^1/2^, **H**/*M*) and **Z** = √*n* **RB** + **E E** ∼ MN(0,**R**,*C*), with *n* = 3 × 10^5^ per trait and *M* = 2 × 10^5^ (per-SNP signal *nh*^2^/*M* ; *h_a_*^2^ ∼ Beta(2, 8)). Scenario S1 (main): two regions on different chromosomes, each of two AR(1) LD blocks of 100 SNPs with neighbour correlation 0.3, 0.6 or 0.9; enrichment *w* non-zero for 10% of SNPs (log-normal, *σ* = 1, mean 1); three cross-region causal pairs with correlations (*ρ*, *ρ*, −*ρ*), *ρ* ∈ {0, 0.5, 0.8}; *q* ∈ {30, 100, 300}; traits either independent (*C* = *I*) or in 10 clusters with within-cluster *r_g_* = 0.6 and full sample overlap with phenotypic correlation 0.3 within clusters. Candidates were the SNPs with *w* > 0 (oracle prescreen; estimated *^w* in S4). Methods: LRCP-GLS (null and plug-in SEs), LRCP-OLS, naive Pearson *r* with the Fisher test on *q* − 3 df, and the oracle correlation of the true rows of **B**. 200 replicates per cell, seed 42; null pairs are all other candidate pairs (about 116,000 per cell). Further scenarios S2-S9 are listed in Supplementary Table S3.

### Data

UK Biobank GWAS summary statistics from Pan-UKB [Karczewski 2025] (European ancestry, both sexes; the 452 phenotypes passing Pan-UKB QC, of which 150 form Pan-UKB’s maximal independent set). The motivating analysis (Fig. 1) uses all 452; LRCP uses the 444 traits analysed by LRCQ, which excludes eight prescription-medication traits whose summary-statistic files were unavailable. LD from UK Biobank in-sample LD [Weissbrod 2020] and 1000 Genomes EUR [1000 Genomes 2015] in Berisa-Pickrell blocks [Berisa & Pickrell 2016]. Motivating PheWAS from the Common Metabolic Diseases Knowledge Portal and associations from the GWAS Catalog. Validation resource: GO biological process [Gene Ontology Consortium 2023] (org.Hs.eg.db 3.18; sizes 10-500).

Summary statistics for the targeted regions were extracted from the tabix-indexed Pan-UKB files. *Z* = *β*_EUR_/SE_EUR_. *n_a_* is the total sample size (cases plus controls for binary traits), the *n* with which Pan-UKB fitted its observed-scale LDSC heritabilities, so that *n_a_*and *h*^2^ come from one fit. Using the effective sample size instead would understate *s_a_* = *n_a_h*^2^/*M* for binary traits by *N*_eff_ /*N* (median 0.33 over the 226 binary traits). *M* = 1,094,844 is the number of HapMap3 SNPs whose LD enters **R**. It differs from the *M*_5_50_ count that Pan-UKB used with full-panel LD scores, because our LD scores are computed within the HapMap3 set. A direct check supports this scale: in the six targeted regions the observed mean *χ*^2^ signal divided by its expectation under *w* = 1 has median 1.47 across traits with *M* = 1,094,844 (1.43 for binary and 1.56 for quantitative traits with total *n*), against 9.1 with *M*_5_50_ and 3.5 for binary traits with *N*_eff_ . The excess over 1 reflects the enrichment of these regions.

### Replication cohorts

We matched Pan-UKB traits by content to GWAS from two cohorts without UK Biobank participants. FinnGen release 12 [Kurki 2023] contributed 57 traits: 35 clinical laboratory measurements (sample-wise median values from the national laboratory database), height, body-mass index and weight, and 19 disease endpoints (median *n* 376,471; range 26,975-500,348). BioBank Japan [Sakaue 2021] contributed 39 traits from its BBJ-only analyses, 33 quantitative traits and 6 diseases (median *n* 145,502; range 71,221-178,726; GRCh37 files from the GWAS Catalog). Variants were matched to the 1,094,844 HapMap3 SNPs by position and both alleles (FinnGen endpoints by rsID), kept if their allele frequency was in [0.01, 0.99], and their *Z* oriented to the Pan-UKB alternative allele; *n_a_* was the total sample size. In each cohort, stages 1-2 were re-estimated from the matched traits by univariate and cross-trait LDSC with Pan-UKB’s EUR (FinnGen) or EAS (BioBank Japan) LD scores and their *M*_5_50_, and stages 3-4 used the same windows, the same 1,799 candidate tags and the same conditional screen as the discovery analysis, with LD from 1000 Genomes EUR (503 individuals) or EAS (504 individuals) [1000 Genomes 2015]. A tag missing in more than half of the traits was untestable; other missing *Z* were imputed at their expectation under *w* = 1. A discovery pair was counted as replicated by its one-sided *P* in the direction of the discovery *z*, with Benjamini-Hochberg or Bonferroni correction over the testable pairs. For reference, the same procedure was applied to Pan-UKB restricted to the matched traits; this reuses the discovery participants and is not a replication.

### Software

All estimators and tests are implemented in the R package lrcpq (version 0.3.1, https://github.com/unfated/lrcpq), whose releases also include the 1000 Genomes EUR HapMap3 LD reference used here.

### Pathway validation

Tags were mapped to their nearest protein-coding gene, as in the companion LRCQ analysis. A gene pair counts as co-annotated if both genes belong to at least one common GO biological process of 10-500 genes [Gene Ontology Consortium 2023] (annotations including those inherited from child terms, from the Bioconductor org.Hs.eg.db 3.18 package; GO release 2023-07-27). We compared the co-annotated fraction of LRCP links with two nulls, each with 500 draws: random distal pairs of the same candidate tags, and degree-preserving rewiring of the links. We also compared it with the same number of top pairs by naive |Z correlation|.

## Supplementary

Supplementary Note: theory of LRCQ and LRCP (model, moment identity, estimators, variances, gene-level tests and power; §S1-§S9 and Appendix A).

Table S1: most correlated distal gene pairs in gene-level association statistics across 4,756 GWAS Atlas studies.

Table S2: Pan-UKB trait list (452 traits; 150 in the maximal independent set; 444 analysed).

Table S3: simulation scenarios S1-S9, their settings and full results. Table S4: prescreening simulation S4.

Table S5: genome-wide modules (all loci and genes) and gene-level burden tests within the 14 modules (899 gene pairs).

Table S6: small-design simulation S8 (regression with each pair counted once or twice, likelihood and naive correlation).

Table S7: calibration of the genome-wide screen against the simulated null, overall and by decile of genetic share, with calibrated P and q for all pairs with *P* < 10^−3^.

Table S8: power of the genome-wide screen for planted correlations (spike-in).

Table S9: type-I error of naive phenome-wide correlation tests under UK Biobank-like trait correlation and sample overlap.

Table S10: replication of the 437 genome-wide hits in FinnGen R12 and BioBank Japan: matched traits, summary by cohort, and per-pair discovery and replication statistics (with the Pan-UKB matched-trait reference).

Figs. S1-S5, with legends, are in the Supplementary Information.

## Supporting information

Supplementary Information: Supplementary Figures S1-S5 and Supplementary Note (theory of LRCQ and LRCP)

## Acknowledgements

We thank Justin D. Tubbs for contributions to an earlier version of this work.

## Author contributions

L.G.C. conceived the study, developed the method and its theory, implemented the software, performed the simulations and the real-data analyses, and wrote the manuscript. T.Q.T. contributed to the revision and audit of the manuscript and to supplementary real-data analyses. P.C.S. contributed to the theoretical conception of the method, reviewed the data analyses and guided the revision of the manuscript.

## Data and code availability

Pan-UK Biobank GWAS summary statistics and LDSC estimates are available from the Pan-UK Biobank project (https://pan.ukbb.broadinstitute.org). UK Biobank in-sample LD matrices (Weissbrod et al. 2020) are at s3://broad-alkesgroup-ukbb-ld, and the PheWAS data are from the Common Metabolic Diseases Knowledge Portal. FinnGen release 12 summary statistics are available from the FinnGen project (https://www.finngen.fi/en/access_results), and BioBank Japan summary statistics from the GWAS Catalog (GCST90018722-GCST90018759 and the six disease studies listed in Table S10). Supplementary Tables S1-S10 are provided as one Excel workbook. The R package lrcpq (version 0.3.1) is available at https://github.com/unfated/lrcpq; its releases include the 1000 Genomes EUR HapMap3 LD reference. All analysis code will be made publicly available upon journal publication and is available from the corresponding author on request.

